# JAK/STAT signaling promotes the emergence of unique cell states in ulcerative colitis

**DOI:** 10.1101/2024.01.22.576736

**Authors:** Grzegorz Maciag, Stine Lind Hansen, Kata Krizic, Lauge Kellermann, Svetlana Ulyanchenko, Martti Maimets, Astrid Møller Baattrup, Lene Buhl Riis, Konstantin Khodosevich, Toshiro Sato, Ole Haagen Nielsen, Kim B. Jensen

## Abstract

The intestinal epithelium forms a barrier to the lumen and ensures uptake of vital nutrients. During inflammatory diseases, such as ulcerative colitis (UC), this barrier is compromised. Here, we perform single-cell RNA sequencing (scRNA-seq) of epithelial cells and outline patterns of cell fate decisions in healthy individuals and patients with UC. We demonstrate that lineage biased progenitors are major sources of normal tissue replenishment, and that patterns of cell behavior are profoundly altered in UC. We furthermore identify unique regenerative cell states linked to JAK/STAT activation extending into the non-inflamed areas of the colon. In organoid models, this can be mimicked by cytokine mediated activation of JAK/STAT leading to the emergence of cell populations with regenerative potential. These findings have profound implications for our understanding of tissue regeneration and illustrates how cytokine signaling influences cell fate decisions. This suggests widespread consequences in patients with chronic ulcerative conditions.

## INTRODUCTION

The epithelial lining of the colon forms the physical barrier protecting our body against the microbes and food antigens present within the intestinal lumen (Gehart and Clevers, 2019). The epithelium is maintained by intestinal stem cells (ISCs) that reside at the bottom of crypts of Lieberkühn and give rise to proliferating progenitors, which further differentiate into absorptive colonocytes and secretory cells including mucus producing goblet cells and hormone secreting enteroendocrine cells (EECs) (de Sousa and de Sauvage, 2019). Upon damage the epithelial barrier heals rapidly to restore its protective function. Any delay in healing can promote sustained inflammation, which triggers further damage and increases the risk of developing inflammatory bowel disease (IBD), including UC (Hindryckx et al., 2016; Kobayashi et al., 2020). Our understanding of disease etiology and factors influencing UC development and progression is gradually increasing (Graham and Xavier, 2020), however, it remains unclear how cell fate is regulated at sites of injury.

Recent advances in single-cell analysis have provided novel insights into the cellular complexity of the colonic tissue. This includes detailed analysis of colonic mesenchyme during homeostasis and ulceration (Kinchen et al., 2018), identification of a new subtype of epithelial cells in the homeostatic epithelium (Parikh et al., 2019), and mapping potential interactions between epithelial, immune and stromal cells (Corridoni et al., 2020; Smillie et al., 2019). However, the principles governing maintenance of the human colonic epithelium, and how this is affected in UC, are incompletely understood. Most of our knowledge about cellular behavior originates from studies using murine models. These studies support a hierarchical model, where ISCs during homeostasis fuel tissue replenishment (Beumer and Clevers, 2021), whereas upon damage cells deviate significantly from these patterns of cell behavior with evidence of cellular dedifferentiation (Yui et al., 2018). Although, analyses have mapped expression of fetal-like gene signature in IBD tissue (Elmentaite et al., 2020; Fawkner-Corbett et al., 2021), the cell fate changes associated with ulceration in patients are largely unexplored.

Mechanistically, numerous signaling pathways linked with cell state changes have been implicated in tissues damage (Larsen and Jensen, 2021). How the human colonic epithelium responds to damage remains to be resolved at both the cellular and molecular level. To address these remaining questions, we analyzed epithelium from healthy individuals and patients with UC using scRNA-seq and utilized human colonic organoids to model predictions made from the computational analysis. We show that distinct progenitors are the main driving source of tissue renewal during homeostasis, and that normal cell fate decisions are disrupted in UC. Furthermore, we identify unique inflammation associated (IA) cell states in the tissue from patients with UC, which are linked with JAK/STAT activation. Lastly, we provide evidence that cells in this cytokine induced state retain proliferative capacity, and following cytokine withdrawal they can revert to a homeostatic stem cell state. Our data suggests that these unique cell states play a role in the regeneration of the colonic epithelium following tissue damage.

## RESULTS

### Proliferative progenitors are the main source of tissue renewal during homeostasis

The cellular composition of tissues and patterns of cell fate decisions can be inferred from scRNA-seq data (Wagner and Klein, 2020). To characterize the epithelium of the sigmoid colon, we analyzed 11,035 epithelial cells by scRNA-seq from four healthy individuals (Figure 1A; S1A-C; Table S1). Cluster analysis combined with expression of specific cell type markers enabled identification of 13 distinct cell clusters representing ISCs (stem), distinct populations of transit amplifying (TA) cells, goblet cells, EECs, tuft cells, and two types of colonocytes (Figure 1C; Table S2) (Parikh et al., 2019). Trajectory inference based on high-resolution clustering, separated the cells into five major lineages with colonocytes clearly divided into classical and pH sensing BEST4+ colonocytes (Figure 1D). Next, we performed RNA velocity analysis that utilized the relative proportion of unspliced versus spliced counts for individual RNA species (Figure S1D). We observed a strong directionality from the ISC population via distinct TA populations towards fully differentiated cell types along all five lineages (Figure 1E). A TA population, which included cells in S and G2/M phase (Figure S1E), was found at the root of each trajectory. Importantly, in tissue sections the proliferation marker PCNA was detected in both TFF3 positive (goblet cells) and negative cells (Figure 1F), suggesting that cell type specification occurred in proliferative TA-cells. Interestingly, the fraction of cells in the TA clusters in S and G2/M was significantly higher than in the stem cell cluster (Figure S1F), which was also supported by the observation that most dividing cells were located above the ISC zone (Figure S1G-H). Furthermore, we observed cells forming unique clusters at the end of the goblet cells and two colonocyte lineage trajectories. Spatial transcriptomics confirmed that cells expressing markers of these end-point clusters (*CEACAM1* and *KRT20*) were located at the crypt top (CT) and facing the lumen both in *MUC2* (goblet cells) positive and negative cells (Figure 1G-I). RNA transcripts associated with the proliferating cells (*HMGB2*) were detected in the bottom third of crypts, spatially confirming implications of the previous trajectory analysis. Moreover, *MUC2* were observed mainly in the top part of the crypts, nevertheless we also observed *HMGB2* and *MUC2* double positive cells supporting the model, where discrete TA populations were responsible for tissue maintenance.

**Figure 1.**
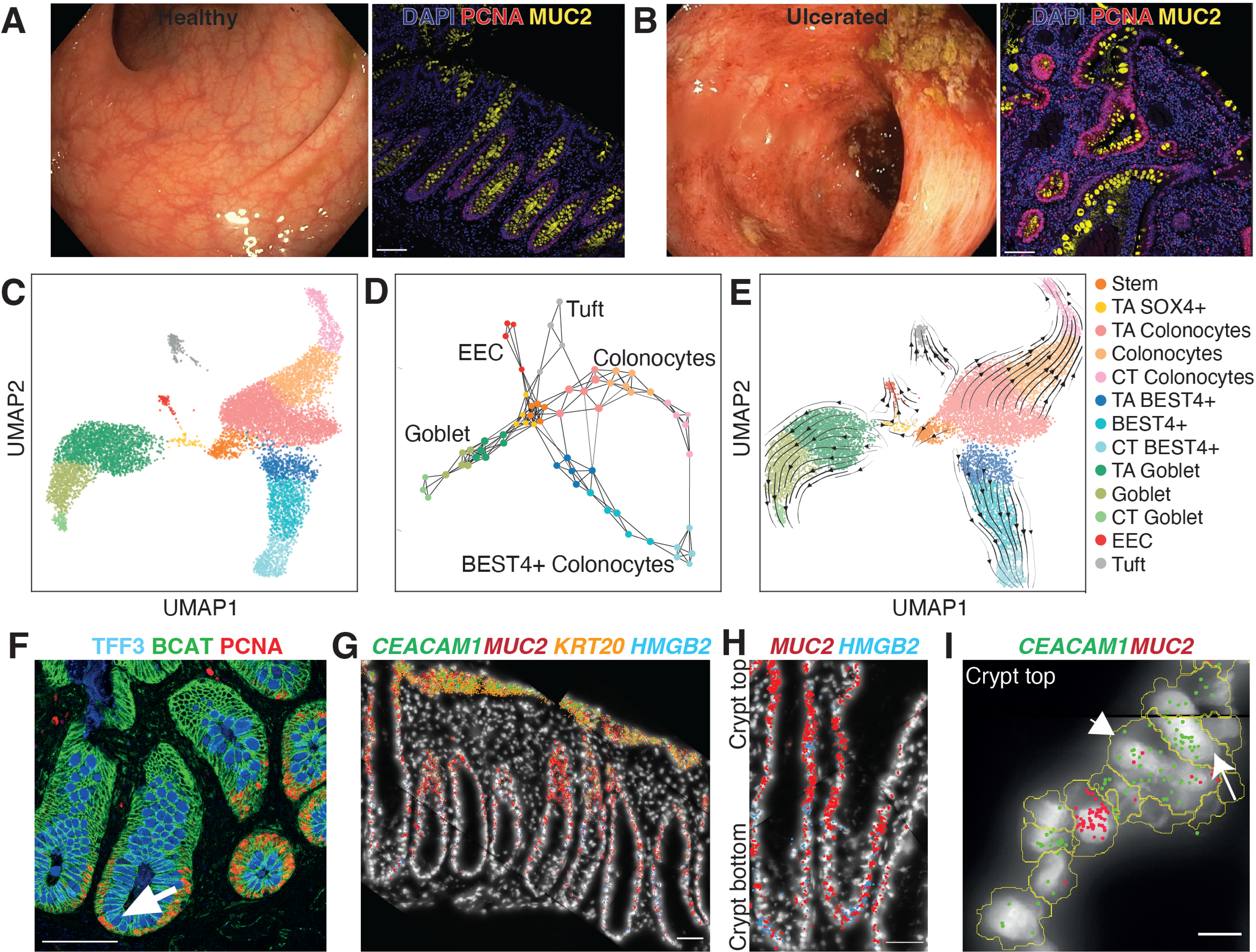
Characterization of healthy human colonic epithelium. **A and B**) Changes in inflamed tissue explored by colonoscopy and immunostaining. A): healthy colon, B): colon from a patient with UC. Goblet cells are marked with MUC2 (yellow) and proliferating cells with PCNA (red), counterstained with DAPI. Scale bar: 100 µm. **C)** UMAP of 11,035 healthy colonic epithelial cells annotated based on marker expression. **D)** Cell type trajectories based on over-clustering. Nodes represent clusters colored by cell type of origin, while edges indicate high degree of similarities. **E)** RNA velocity measurements projected onto the UMAP embedding shown in (B). **F)** Immunostaining for TFF3 (blue), PCNA (red) and β-Catenin (green) in healthy colonic tissue. A proliferating goblet cell is indicated with an arrow. Scale-bar: 100 µm. **G, H and I)** Spatial distribution of RNA molecules for *CEACAM1* (green), *MUC2* (red), *KRT20* (orange) and *HMGB2* (blue), counterstained with DAPI. F and G show entire crypts. H presents a zoom of a representative crypt-top area. CT colonocyte (arrow) and goblet cell (arrowhead) are indicated. Scale-bar: 50 µm (F and G) and 10 µm (H). See also Figure S1

### Ulceration promotes changes in cell behavior

To outline how the patterns of cell fate decisions were affected in patients with UC, we delineated cellular relationships in disease affected tissues. We obtained biopsies from visibly inflamed parts and the neighboring healthy margin of the sigmoid colon. Tissue damage in the ulcerated regions was evident at the morphological level, with crypts undergoing remodeling and lacking a defined crypt top layer, while epithelium from the healthy margin adjacent to the inflamed areas were morphological similar to healthy tissue (Figure 1B, 2A). At the cellular level the inflamed areas were associated with pronounced changes affecting cellular composition with a higher proportion of CD45+ cells and a lower fraction of epithelial cells, when compared to the homeostatic tissue, and the healthy margin of the same patient (Figure S1B). To generate a map reflecting the cellular heterogeneity in patients with UC, we performed scRNAseq on 4,351 epithelial cells from inflamed area and 6,762 cells from healthy margin from the same patients with UC, identifying largely the same cell populations as observed in healthy tissues (Figure 2B; S2A). Thus, the cell type analysis revealed that the epithelium from healthy margin and the inflamed areas of UC patients contained largely the same cell populations found in the homeostatic epithelium.

**Figure 2.**
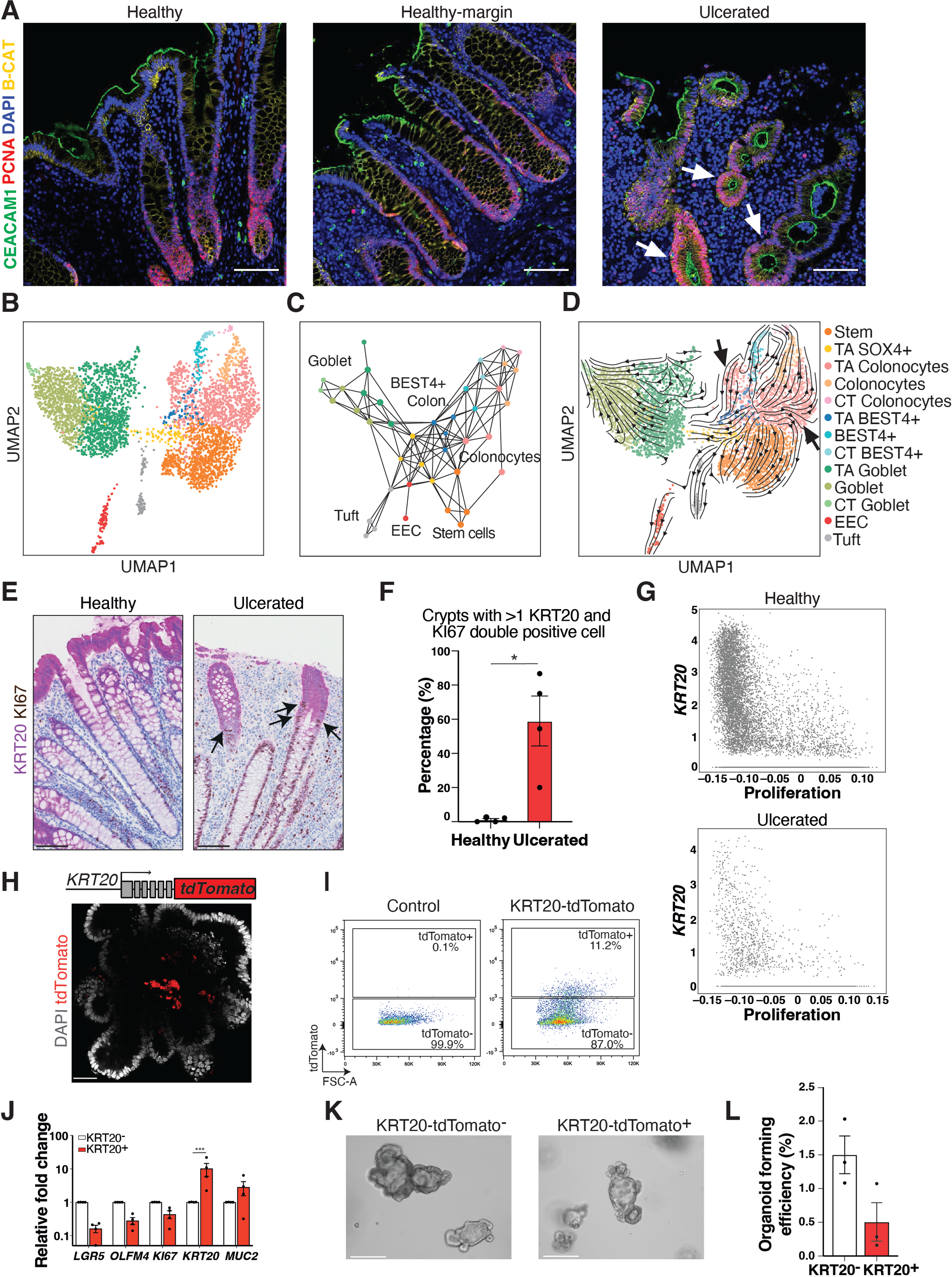
Pronounced lineage infidelity in tissue regeneration. **A**) Immunostaining for CEACAM1 (green), PCNA (red) and β-catenin (yellow) in healthy, healthy-margin and ulcerated colonic tissue. Scale-bar: 100 µm. **B)** UMAP depicting 4,351 colonic epithelial cells from patients with UC annotated as in the healthy dataset. **C and D)** Trajectory graph based on over-clustering and RNA velocity. Arrows (D) indicate trajectories going from differentiated cell clusters towards progenitor clusters. **E)** Representative IHC for KRT20 and KI67 of healthy and ulcerated human colon. Arrows indicate KRT20 and Ki67 double positive cells. Scale bar: 100 µm. **F)** Percentage of crypts with >1 double KRT20/KI67-positive cells in healthy and ulcerated human colon. Data shown as ±SEM; n=4 patients; \**P*<.05 two-tailed unpaired *t*-test. **G)** Scatter plots of *KRT20* expression (y-axis) and proliferation score (x-axis) in the healthy (top) and ulcerated (bottom) datasets. **H)** Representative image of tdTomato (red) and DAPI (grey) in the KRT20-tdTomato human colon organoid reporter. Scale bar: 50 µm. **I)** Flow cytometry analysis for tdTomato in the KRT20-tdTomato human organoid reporter model. **J)** Expression levels of indicated genes of KRT20-tdTomato^-^ sorted organoids (white) and KRT20-tdTomato^+^ sorted organoids (red) normalized to *GAPDH* expression. Fold change relative to KRT20-tdTomato^-^ cells ± SEM; n=4 individual experiments; \*\*\**P*=.0004 ordinary 2way AVOVA with Sidak’s correction for multiple comparison. **K)** Representative images of human colon organoids formed from tdTomato^-^ (top) and tdTomato^+^ (bottom) cells 10 days following seeding. Scale bars: 275 µm. **L)** Quantification of organoid-forming efficiency of KRT20-tdTomato^-^ and tdTomato^+^ cells. Data shown as mean ±SEM; n=3 independent experiments with average of four wells counted for each experiment with 2,500 cells seeded per individual well. See also Figure S2

A deeper analysis of the dataset from the ulcerated regions revealed pronounced changes. Trajectory analysis showed striking differences with shorter branches along the three major cell lineages suggesting that the common programs of epithelial differentiation were disrupted (Figure 2C). Additionally, RNA velocity analysis indicated that normal differentiation patterns in the dataset from healthy individuals were altered in the ulcerated samples (Figure 2D). Notably, trajectories now existed into and out of the stem cell compartment (Figure 2C-D), while normal patterns of differentiation were observed in the healthy margin data set (Figure S2B-C). Keratin 20 (KRT20) is a known marker of terminally differentiated cells in the colon and can be found in the CT populations across the datasets (Figure S2D). In line with the observed changes in the cellular trajectories in the diseased samples, we could detect cells co-expressing the differentiation marker KRT20 and KI67, a marker of proliferating cells, at the protein level in more than 50% of the crypts analyzed, suggesting that this was a recurrent event unique for patients with UC (Figure 2E-F; S2E). Interestingly, we did not detect such co-expressing cells in the single cell data, and cells with high *KRT20* expression generally had a very low proliferation scores, not only in healthy, but also in the diseased epithelium (Figure 2G). Given the long half-lives of keratins(Denk et al., 1987), the detection of KRT20 and KI67 double positive cells suggested that cells that previously expressed *KRT20* had reentered the cell cycle. Collectively, this demonstrated that ulceration alters the normal patterns of cell behavior.

### Human colonic epithelial cells expressing the differentiation marker KRT20 display self-renewal potential

Based on the detection of KRT20 and KI67 double positive cells in inflamed areas of UC epithelium, we hypothesized that KRT20 expressing cells in a human context retained the ability to self-renew, when placed in a growth permissive environment. To test this hypothesis, we utilized a human organoid model where tdTomato was introduced into the 3’UTR of the *KRT20* locus (Figure 2H) and purified tdTomato positive and negative cells (Figure 2I). As expected tdTomato positive (KRT20^+^) cells showed elevated levels of differentiation markers (*KRT20* and *MUC2*), and reduced expression of markers of stem cells (*LGR5* and *OLFM4*) and proliferation (*KI67*), when compared to tdTomato negative (KRT20^-^) cells (Figure 2J). The self-renewal potential of tdTomato positive and negative cells was subsequently tested using organoid formation assays. As expected, tdTomato negative (KRT20^-^) cells readily formed budding organoids, however, supporting our findings from the patient data, a proportion of tdTomato positive (KRT20^+^) cells likewise retained this capacity, although to a lesser extend (Figure 2K-L). This supported that cells expressing markers of differentiation can return to a proliferating state, when placed in growth-permissive environment.

### Emergence of unique cell populations following ulceration

Based on the findings that the cellular trajectory was altered during inflammation, we next integrated the three scRNAseq data sets (healthy, ulcerated and healthy margin) to directly investigate changes in the cellular complexity. Cluster analysis revealed that the overall patterns of cell populations followed the cell types detected in the healthy colon, however, with changes in the relative distribution of cells within the distinct cellular clusters (Figure 3A-C). Firstly, it was clear that some of the observed changes were observed both in the ulcerated regions and the neighboring healthy margins. Notably, changes could also be observed in the tissue where CD74, an immune cells marker for e.g., B lymphocytes, macrophages, dendritic cells, could be detected in the mucosa in both the ulcerated regions and the neighboring healthy margin suggesting an influence of systemic inflammation during disease (Figure S3A). Moreover, *CD74* could also be detected at the RNA level in the scRNAseq datasets, with higher proportions in the samples from UC patients. Using CD74 to subdivide the cell populations in the samples from the healthy margin revealed that CD74 expressing cells based on Pearson correlations of the first 50 principal components were more representative of the ulcerated data-set, whereas CD74-cells represented an intermediate between the ulcerative and healthy state. Secondly, samples from the ulcerated and the healthy margin had an increased proportion of cells expressing stem cell markers (Figure 3C). Notably, there is not only a larger fraction of stem cells, but cells within the stem cell cluster were transcriptionally distinct from their healthy counterparts (Ulcerated vs Healthy: 881 up and 1063 down regulated genes; Healthy-margin vs Healthy: 284 up and 203 down regulated genes; Log2FC>+/-0.5, p<0.05; Figure S3B). Moreover, there was a significant overlap in the genes that were detected as differentially expressed in the healthy-margin and ulcerated samples, when compared to the healthy samples (Up: p=9×10^-158^; Down p=6×10^-185^). Here genes upregulated in the ulcerated and non-ulcerated stem cell compartment revealed enrichment for genes associated with interferon signalling and antigen processing, whereas genes associated with metabolic processes were repressed (Figure S3B). This supported the absence of a sharp border between affected and non-affected tissues. Thirdly, two new cell populations within the colonocyte and goblet cell branches appeared in the ulcerated samples (Figure 3A-C). The two new cell populations were only detected in epithelium of UC patients and were accordingly named inflammation associated (IA) colonocytes and IA goblet cells. Cells in the IA clusters were characterized by a high proportion of proliferating cells (Figure 3D), and clustered independently in the ulcerated dataset (Figure 3E). GO-term analysis for the differentially expressed genes in both the IA goblet and colonocyte cell populations, when compared to the rest of the cell clusters, showed enrichment of cytokine and interferon signaling, immune response and antigen processing, similar to what was observed for the stem cell compartments (Figure 3F-G). The data therefore suggested that these new IA clusters contained cells in a proliferative state that were linked with signals originating from the immune system.

**Figure 3.**
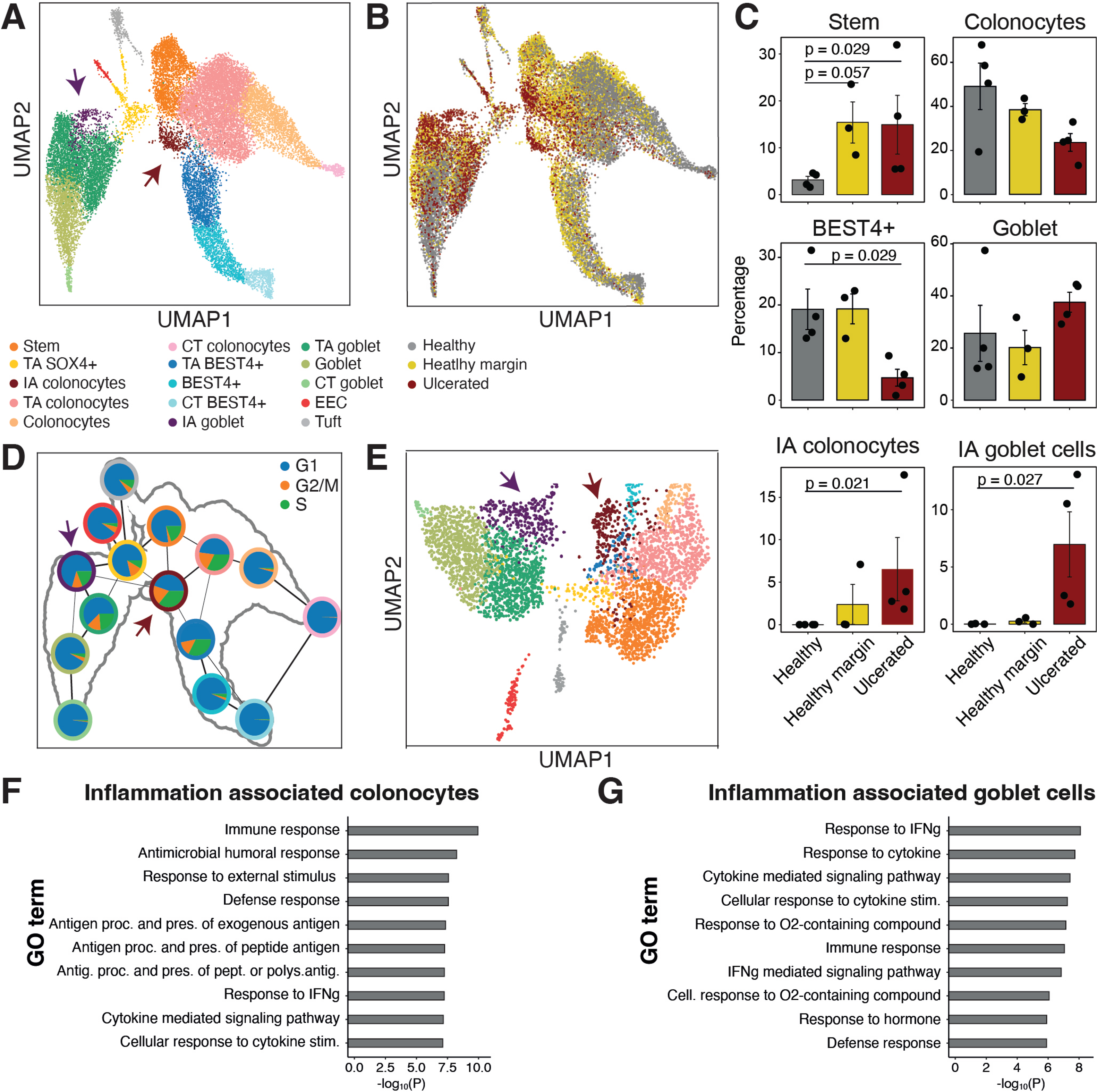
Emergence of new cell states in ulcerated tissues. **A and B**) UMAP of the combined datasets from healthy, healthy margin and ulcerated epithelial cells colored by cell type cluster and dataset of origin. Arrows indicate the IA cell states. **C)** Cell fractions representing the major cell lineages in the healthy, healthy margin and ulcerated datasets, including the IA cell states. Colonocytes, BEST4+ and goblet cell lineages include TA and CT populations. Significance tested with two-sided Mann–Whitney *U* test. Error bars represent the S.E.M. **D)** Trajectory graph of cell type clusters in the integrated datasets. Pie charts indicate proportions of cells in each cell cycle phase. Outer ring colors matches cell types in (A). **E)** UMAP with cell type clusters in the ulcerated dataset including annotation of new IA cell states (arrows). **F and G)** Top-10 GO terms from overrepresentation analysis of differentially expressed genes calculated by comparing expression of genes in a given cluster to the rest of the dataset. See also Figure S3

A number of key pathways influence cell behavior following tissue damage and promote tissue regeneration. To infer which gene regulatory networks were associated with the emergent IA cell populations we utilized DoRothEA (Garcia-Alonso et al., 2019). Aligned with the enrichment of GO-terms associated with both cytokine and interferon signaling the most enriched regulatory network at the single cell and population level was STAT3 signaling (Figure 3F-G; 4A; S3C-D; Table S3; p<10^-100^). To further investigate the features of the new cell states, we examined genes uniquely expressed by the two IA populations compared to the lineage they were closest to (goblet cell and colonocytes). Less than 100 genes were found to be differentially expressed in each population, including several reported STAT target genes observed in both cell states (*REG1A*, *REG1B*, *REG3A*, *DMBT1*, *ALDH1A1*, *C4BPB* and *IFITM1*; Figure 4B). The analysis identified *REG1A (*Regenerating Family Member 1 alpha) as a marker unique to the two IA cell states (Figure 4C). Examination of independent datasets confirmed the expression of *REG1A* across samples from patients with UC (Figure S3E). At the cellular level, REG1A was detected in ulcerated epithelium in both TFF3 positive and negative cells, supporting the emergence of these cell states in the goblet cell and colonocyte lineages (Figure 4D).

**Figure 4.**
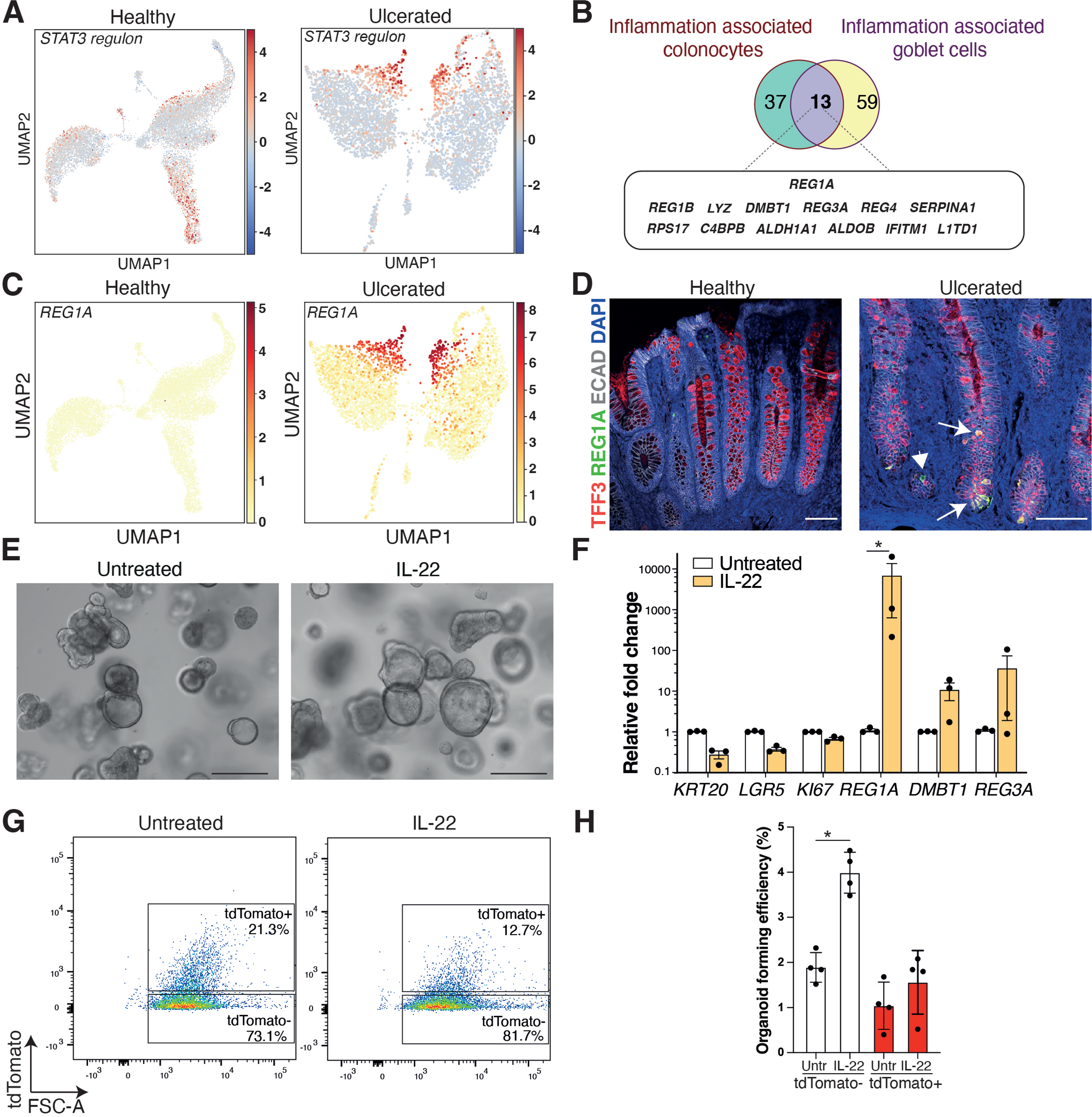
JAK/STAT signaling is linked with the appearance of inflammation associated cell states. **A**) UMAP with STAT3 regulon score in healthy and ulcerated datasets. **B)** Venn diagram of differentially expressed genes in the emergent IA cell populations. **C)** UMAP with a heatmap of normalized *REG1A* expression in healthy and ulcerated datasets. **D)** Immunostaining for REG1A (green), TFF3 (red), and E-Cadherin (white) in healthy and ulcerated tissue. Arrows indicate double positive cells, while an arrowhead indicates REG1A positive, but TFF3 negative cells. Scale bars: 100 µm. **E)** Bright field images of control and IL-22 treated human colonic organoids. Scale bars: 275 µm. **F)** Expression levels of indicated genes of control (white) and IL-22 treated organoids (yellow) normalized to *GAPDH* expression. Fold change relative to control treated cells ±SEM; n=3 independent experiments; \**P*<.05 ordinary 2way ANOVA Fisher’s LSD test. **G)** Representative flow cytometry plots of colonic epithelial cells isolated from control and IL-22 treated human colon KRT20-tdTomato organoids. **H)** Quantification of organoid forming efficiency of KRT20-tdTomato^-^ and tdTomato^+^ cells from control or IL-22 treated organoids. Data shown as mean ±SD; n=4 independent experiments for each population with 2,500 cells initially seeded per well; * *P*<.05 unpaired Mann-Whitney t test. See also Figure S3 and S4

Thus, cells in de novo cell states in UC were characterized by being actively cycling, expressing a unique gene signature associated with cytokine signaling and STAT pathway activation.

### JAK/STAT activation promotes cell state changes

The JAK/STAT signaling pathway mediates inflammatory and proliferative responses downstream of cytokine stimulation. Interleukin-22 (IL-22) is one of several cytokines that activates STAT signaling during tissue regeneration and is known to be involved in tissue-repair following intestinal damage in murine models (Cox et al., 2021; Lindemans et al., 2015). To investigate how STAT activation affected human colonic epithelium, we therefore treated colonic organoids from healthy individuals with IL-22 (Figure 4E). The treatment led to a reduction in expression of markers associated with differentiation (*KRT20*) and stem cells (*LGR5*), as well as upregulation of genes associated with the emergent IA cell states identified in patient samples (*REG1A, DMBT1* and *REG3A*; Figure 4F). The induction was JAK dependent, and not affected by mTOR inhibition, which previously has been shown to support IL-22 induced Paneth cell differentiation (Figure S3F-J) (He et al., 2022). Importantly, IL-22 treated cells retained the capacity to grow (Figure 4E-F), and when grown in the absence of R-Spondin1 and WNT, IL-22 treatment reduced the proportion of Krt20-tdTomato+ cells (Figure 4G) and enhanced the organoid forming capacity specifically of the Krt20-tdTomato-population (Figure 4H).

To assess the effect of inhibiting JAK/STAT signaling during tissue repair *in vivo,* we assessed the impact of pathway inhibition on tissue regeneration following the treatment of animals with dextran sulfate sodium (DSS). Here mice were treated with the pan-JAK inhibitor, tofacitinib, during the early repair phase. It was evident that tofacitinib treated animals developed more severe colonic damage and greater weight loss when compared to control animals (Figure S4A-C). Thus, as described for STAT3 (Oshima et al., 2019), JAK/STAT signaling was required for initiating epithelial regeneration in vivo suggesting that the benefits observed in patients arose from the effect of pan-JAK inhibition on the inflammatory process.

Collectively, this demonstrated that cytokine signaling was essential for tissue regeneration, that IL-22 promoted changes that mirror what was observed during tissue regeneration in vivo and that IL-22 promoted self-renewal under non-permissive conditions. Taken together, these observations demonstrated that intricate signaling relationships between epithelial and immune cells were important for cellular responses following damage.

### Cytokine induced JAK/STAT signaling is sufficient to promote the IA cell state

To address whether cytokine signaling was sufficient to promote the emergence of the new IA cell state, we performed scRNA-seq on organoids from 3 healthy individuals cultured with or without IL-22 (Figure 5A). In control samples, all major cell populations except for BEST4+ colonocytes could be identified, whereas the treated samples included cell clusters transcriptionally matching the novel IA states observed in patients with UC (Figure 5B-C). Similar to the patient data, these new clusters appeared in the goblet cell and colonocyte, were unique to the IL-22 treated organoids and showed *REG1A* expression and STAT pathway activation (Figure 5D-E; S4D-E).

**Figure 5.**
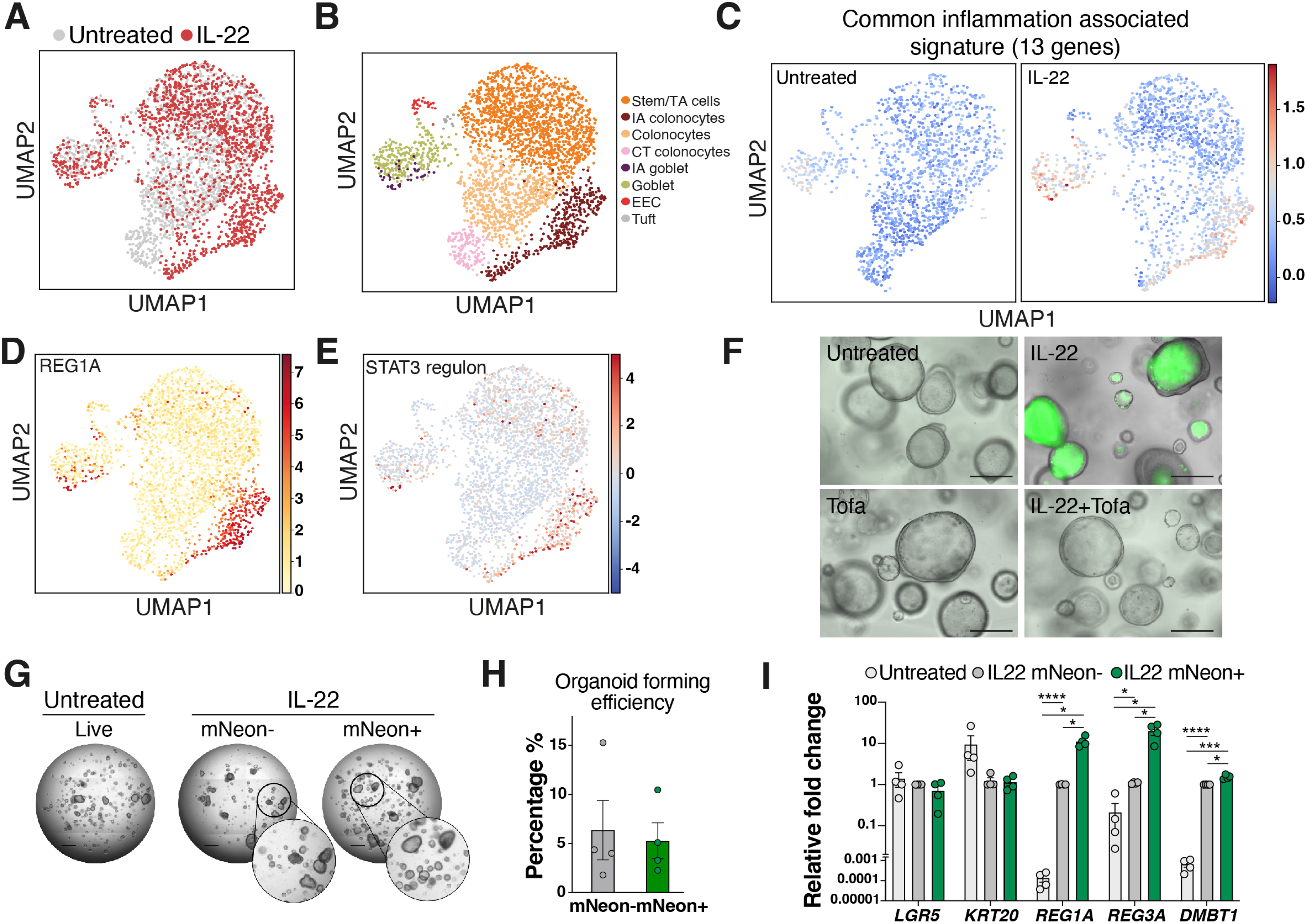
The IA states are induced in human organoids through JAK/STAT activation. **A and B**) UMAP of untreated and IL-22 treated organoid datasets colored by condition and cell type from organoids derived from 3 different individuals. **C)** UMAP with a heatmap of average expression of the common emergent signature (Figure 4B) in control (left) and IL-22 treated (right) organoids. **D and E)** UMAP with heatmaps of normalized *REG1A* expression and STAT3 regulon score in the organoid dataset. **F)** REG1A-mNeonGreen tagged human organoids untreated or treated with IL-22 and/or tofacitinib (Tofa) for three days. Scale bars: 275 µm. **G)** Representative images of REG1A-mNeonGreen tagged human organoids re-seeded in Matrigel after purification through flow cytometry, either untreated (live cells) or treated with IL-22 for three days (mNeonGreen+ and mNeonGreen-cells). Scale bars: 1mm. **H)** Quantification of organoid forming efficiency related to G. Data shown as ± SEM; n=4 independent experiments for each population with 2,500 cells seeded per well. **I)** Expression levels of indicated genes of cells purified through flow cytometry from REG1A-mNeonGreen control cells (light grey) and IL-22 treated cells, either mNeonGreen-(dark grey) or mNeonGreen+ cells (green), normalized to *GAPDH* expression. Fold change relative to mNeonGreen-cells ± SEM; n=4 independent experiments for each population. *p<.05, **p<.01, ***p<.001, ****p<.0001; two-way ANOVA with Tukey’s multiple comparison test. See also Figure S5

Finally, to assess the regenerative capacity of the REG1A population, we engineered a reporter line fusing mNeonGreen to the carboxy-terminus of REG1A (Figure S5A). Upon IL-22 treatment, approximately 15% of the cells expressed REG1A-mNeonGreen, similar proportions to what was observed in the scRNA-seq dataset with accumulation of secreted REG1A-mNeon fusion protein in the lumen of the organoids (Figure S4D, S5B-C). Treatment with JAK or mTOR inhibitors further confirmed that REG1A expression was regulated through JAK/STAT pathway (Figures 5F; S5D-E). Importantly, cells expressing *REG1A* in the scRNAseq dataset were distinct from the population classified as transit amplifying (TA) or stem cells, leaving it unclear whether these cells would have reduced self-renewal capacity compared to the rest of the cell population. To characterize the properties of the REG1A expressing cells, we therefore purified mNeonGreen negative and positive cells by flow cytometry. In self-renewal assays, REG1A-mNeonGreen-negative and –positive cells both formed organoids at a similar efficiency, demonstrating that the cytokine induced REG1A cells retained the capacity to self-renewal and therefore were likely to have the capacity to contribute to regeneration (Figure 5G-H; S4F-G). As previously observed, IL-22 treatment led to reduced expression of *KRT20* and induction of markers of the cytokine-induced state (*REG1A*, *REG3A* and *DMBT1*), most prominently in the REG1A-mNeonGreen^+^ cells (Figure 5I; S5H).

Collectively, our data demonstrated that the emergent cell populations could be recapitulated *in vitro* using organoid models, which enabled us to reveal the *in vitro* regenerative capacity of these novel cell populations.

## DISCUSSION

Here, we present a comprehensive analysis of the human colonic epithelium during steady state homeostasis and in patients with UC. The analysis revealed that during homeostasis early lineage choices were linked with lineage biased TA populations as the main source of tissue renewal. This is in line with recent reports suggesting a minor role of colonic stem cells in the daily tissue replenishment (Ishikawa et al., 2022; Lee-Six et al., 2019; Nicholson et al., 2018). Aligned with studies of mouse models (Denk et al., 1987; Tetteh et al., 2016; Tomic et al., 2018; van Es et al., 2012), the unidirectional flow of cells from the TA compartments was, however, perturbed in patients suffering from ulcerations (Harnack et al., 2019; Tetteh et al., 2016; Tomic et al., 2018; van Es et al., 2012). Using genetically engineered human organoid lines we could show that cells expressing markers of differentiation retained the capacity to self-renew, when provided with a growth permissive environment. It is consequently important to understand how cell types are influenced by the environment and via signaling crosstalk.

Experimental injury models have revealed that the regenerative responses are characterized by loss of markers associated with both differentiated cells and stem cells and the emergence of cells with new properties (Nusse et al., 2018; Yui et al., 2018). Interestingly, in clinical samples we observed that this was recapitulated in both the colonocyte and goblet cell lineages with the emergence of new cell populations with a proliferative potential. Importantly, the emergence of cells with these *in vivo* properties can be recapitulated *in vitro* upon stimulation with a single cytokine enabling us to demonstrate their self-renewal properties. These cell populations are characterized by a strong enrichment of STAT target genes, a feature that has previously been linked to tissue regeneration (Pickert et al., 2009; Taniguchi et al., 2015). Aligned with studies in mouse, it is interesting that *Ly6a* (Sca1), another STAT target gene, is associated with the emergence of cells required for regeneration in the mouse (Rao et al., 2015). IL-22 is one of several secreted factors that activate JAK/STAT signaling and known to be involved in tissue regeneration (Lindemans et al., 2015; Pickert et al., 2009). Notably, blocking JAK signaling in experimental colitis models impaired mucosal healing. Our data combined suggests that in addition to promoting the inflammatory process JAK signaling is similarly important for elements of the epithelial repair. Interestingly, the clinical use of pan-JAK inhibitors has been associated with a wide range of adverse effects like herpes zoster infection, major adverse cardiovascular events (Ytterberg et al., 2022), and more selective JAK inhibitors are now marketed and in the pipeline (Danese et al., 2022; Granau et al., 2023; Nielsen et al., 2023). This highlights that tissue regeneration in humans is associated with pronounced changes in cellular identity and that this is driven via crosstalk between immune and epithelial cells.

Collectively, we have investigated cell behavior in healthy and ulcerated colonic epithelium at high resolution using scRNA-seq combined with organoid models. A deeper understanding of general mechanisms that regulate tissue regeneration will be important for the development of targeted treatment of patients suffering from UC. Studies using mouse models have implicated WNT and YAP signaling in the process of mucosal healing following tissue damage (Gregorieff et al., 2015; Harnack et al., 2019; Yui et al., 2018). Whether these signaling pathways, in combination with JAK/STAT pathways are implicated in tissue regeneration in humans, remain an open question. From a therapeutic perspective, identifying how to promote the induction of regeneration will launch possibilities for more rapid mucosal healing in patients with UC and will potentially be able to complement current treatment strategies.

## EXPERIMENTAL PROCEDURES

### Resource availability

Further information and requests for resources and reagents should be directed to and will be fulfilled by Kim Bak Jensen; a completed Materials Transfer Agreement may be required. Single-cell RNA-seq data generated for this paper have been deposited at the European Genome-phenome Archive (EGA) repository and are publicly available as of the date of publication under the accession number EGAS00001007098. For reviewers, please contact the editorial office, who will provide you with access to the data via EGA.

### Human samples

For single-cell RNA sequencing analysis four patients with an established diagnosis of UC and four controls were included (Patient details are included in Supplementary Table 2). All subjects underwent a routine sigmoidoscopy or colonoscopy as part of their clinical evaluation at the Endoscopy Unit, Department of Gastroenterology, Herlev Hospital due to their clinical symptoms. Biopsies were collected from a visibly inflamed area of the sigmoid colon, and biopsies from an endoscopically normal part of the descending colon were collected at the same time. Controls were recruited from the Danish National Screening Program for Colorectal Cancer where all investigations turned out normal. UC was diagnosed according to well-established criteria (Kobayashi et al., 2020) and included UC patients had an endoscopic Mayo score of either I or II at time of enrolment (Schroeder et al., 1987). Exclusion criteria were as follows: age >80 or <18 years, clinical evidence of infection, use of antibiotics or probiotics within 14 days, severe mental illness, and pregnancy. Healthy control samples were obtained from the national screening program for colorectal cancer where all investigations turned out normal.

For immunohistochemistry and immunofluorescence analysis, formalin fixated and paraffin embedded sections from macroscopic inflamed and non-inflamed colonic areas were collected from surgical specimens of colectomies from four patients with UC after the specimen had undergone evaluation at the Department of Pathology, Herlev Hospital.

Human colon organoid lines were established from the colonic epithelium of three healthy donors and were used to study for the cytokine treatment.

The human colon organoid line genetically modified to report KRT20 expression, KRT20-tdTomato, was derived from a healthy individual and used in previous studies (Shimokawa et al., 2017; Sugimoto et al., 2018).

The study was approved by the Scientific Ethics Committee of the Capital Region of Denmark, filed under the ethical permission number H-18005342. All individuals provided written informed consent to participate in this study. Participants were informed both orally and in writing in compliance with the Declaration of Helsinki and the guidelines of the Danish National Scientific Ethics Committee.

Single-cell RNA-seq data generated for this paper have been deposited at the European Genome-phenome Archive (EGA) repository and are publicly available as of the date of publication under the accession number EGAS00001007098.

### Mice

Wild-type female C57BL/6J mice were obtained from Taconic Denmark. Mice used for this study were between 8-to 15-week and they were housed under conventional conditions and in accordance with institutional regulations in a specific-pathogen-free (SPF) Animal Facility in individually ventilated cages, under a 12hr light-dark cycle. Food and water were provided ad libitum. None of the animals used in these studies had been subjected to prior procedures and were drug– and test-naive.

All animal experiments and procedures were conducted in accordance with the recommendations of the European Community Directive (2010/63/EU) and the project was approved by the Danish Animal Experiments Inspectorate under the license number 2017-15-0201-01381.

### Cell isolation and flow cytometry

Biopsies were washed in cold PBS supplemented with 0.5 mM DTT two-three times before they were incubated in 8 mM EDTA with 0.5 mM DTT in PBS on ice for one hour on a rocking platform. Supernatants were removed and 5 mL cold PBS was added. The tubes were shaken forcefully by hand to free epithelial crypts. The crypt containing supernatants were collected in a 15 mL falcon tube and the shaking was repeated twice to obtain a total volume of 15 mL PBS solution containing crypts. The crypts were spun down at 300 *g* for 4 minutes and the pellets were resuspended in 3 mL TrypLE supplemented with DNaseI (2.5 μg/mL), rock inhibitor Y-27632 (10 μM) and N-acetylcysteine (1 mM) and incubated at 37°C for 10-20 minutes. At the end of the incubation the single cell suspensions was supplemented with 10 mL of advanced DMEM/F-12 (Gibco™) media with 10% adult bovine serum. The single cells were pelleted by centrifugation at 400 *g* for 4 minutes and the pellet was resuspended in 2 mL 1% BSA/PBS with gentle pipetting and pelleted before resuspension in 1% BSA/PBS filtered firstly through 70 μm filter and then through a 40 μm filter. The cells were stained in 200 μL 1% BSA/PBS with EPCAM-APC, CD31-APC-780, and CD45-PE-CF594 on ice for 25 minutes, washed three times with 0.1% BSA/PBS before they were incubated with DAPI (Sigma-Aldrich) in 200 μL for 5-10 min on ice and flow sorted on an ARIA I or ARIA III BD sorters using a 70 μm nozzle. Cells were sorted into an Eppendorf tube with 2 μL ultra clean 0.1% BSA/PBS for the generation of scRNA-seq libraries.

### Single cell library preparation and sequencing

Single cell libraries were prepared using the 10X Genomics protocols v2 Chemistry. Maximum 25,000 sorting events were loaded per well in a volume of 33.8 μL ultra clean 0.1% BSA/PBS. Cells were encapsulated in droplets of Gel Bead-in-Emulsions (GEMs) using the 10X Genomic Single Cell Chip whereafter reverse transcriptase was performed. Purified cDNA was amplified with 10-14 PCR cycles dependent on the number of cells loaded. Libraries were diluted to 2 nM in elution buffer and two libraries were pooled and diluted to 1.7 pM before running on the same flow cell. The libraries were sequenced on an Illumina NextSeq 500 platform with a High Output 150 cycles kit.

### Processing of raw scRNA-seq reads

Cell Ranger (v3.0.1) software from 10× Genomics was used to process Chromium single-cell RNA-seq output to align reads and generate bam files needed for the next steps (Zheng et al., 2017). The refdata-cellranger-GRCh38-3.0.0 reference was downloaded from the 10× Genomics website (https://support.10xgenomics.com/). First, cell ranger mkfastq function was used to demultiplex raw base call files into FASTQ files. Then, the FASTQ files were aligned and filtered with cell ranger count function. One of the outputs of that function were bam files, which were then processed with velocyto using the run10x function (La Manno et al., 2018). Velocyto counts reads falling into exonic/intronic regions and generates spliced/unspliced expression matrices in a loom file.

### Quality control and filtering of cells

The loom files were processed and analysed in Python using scanpy and scVelo (Bergen et al., 2020; Wolf et al., 2018). Low quality cells were filtered out with each sample being processed separately in the following manner. Firstly, per cell distributions of spliced counts, unspliced counts and genes were inspected. Based on visual tracing of a Gaussian curve, the cut-off values for those three characteristics were determined (specific to each sample). Cells below the thresholds for unspliced counts and genes and below or above thresholds for spliced counts were filtered out. Next, cells with a high ratio of counts originating from mitochondrial features were eliminated based on individual ratio distributions (0.3 in most cases, 0.4 and 0.45 in two ulcerated samples). Finally, doublets (two cells in a single bead) were filtered out with both scrublet and DoubletFinder (McGinnis et al., 2019; Wolock et al., 2019). The threshold score for expected doublet rate was set to 0.06 for scrublet, and default parameters were used for DoubletFinder. Cells predicted to be doublets by either of the tools were filtered out.

### Data integration, normalization, and batch correction

After cell filtering, samples for each of the datasets were concatenated together. At this stage, genes expressed in less than 5 cells or with less than 20 counts were filtered out. Count data was then normalized, and log transformed for each cell by total counts over all genes. The top 2000 highly variable genes were used for batch correction using the mutual nearest neighbour’s algorithm implemented as mnn_correct function in scanpy’s API. Parameter k was set to 15 and var_adj to True.

### Cell cycle annotation

Cell cycle stage annotation was performed before batch correction using the score_genes_cell_cycle function from scanpy, based on published cell-cycle signatures (Tirosh et al., 2016).

### Proliferation score calculation

Proliferation score was calculated before batch correction using the score_genes function from scanpy, based on a published proliferation signature (Merlos-Suarez et al., 2011).

### Regression, dimensionality reduction and embedding

Unwanted variation coming from the cell cycle stage was reduced by linearly regressing the annotated S and G2/M scores. After those changes, the top 2000 highly variable genes were selected again for the rest of the analysis. Dimensionality reduction was first performed using principal-component analysis (PCA) and then with Uniform Manifold Approximation and Projection (UMAP). UMAP requires calculation of neighbours for each of the cells, which was done with batch balanced k-nearest neighbours algorithm implemented in the bbknn package (Polanski et al., 2020). The Euclidean metric, 30 principal components and 5 neighbours within batch were used and the parameters approx and use_faiss were set to False.

### Clustering

Initial clustering was performed using the unsupervised Leiden method with a high-resolution parameter, giving rise to multiple clusters. At the same time, expression of canonical cell type markers (*LGR5*: Stem cells, *LRMP*: Tuft cells, *BEST4*: BEST4+ colonocytes, *SCGN*: Enteroendocrine cells, *CA1*: Colonocytes, *MUC2*: Goblet cells) combined with results from differential expression analysis were used to guide cluster identities. Subsequently, multiple iterations of merging and dividing the clusters were performed until a fully informative cell type annotation of clusters was achieved.

### Differential expression

Differentially expressed genes (DEGs) were detected with the Wilcoxon rank sum test using scanpy’s rank_genes_groups function. The test was performed comparing expression of genes in each cell type cluster to the rest of the cells in each dataset.

### Trajectory analysis

PAGA was used to infer developmental trajectories in the healthy and ulcerated datasets. It was run on sub-clusters, which were generated by rerunning the clustering algorithm within each of previously annotated cell type clusters. PAGA works by approximating all sub-clusters to single nodes and connecting them with edges based on similarity (Wolf et al., 2019). This similarity is related to the overrepresentation of edges in the KNN graph between cells in the two clusters. The number of edges were adjusted using the threshold parameter to get 2-4 edges per node.

### Spliced/unspliced count ratio

Using the function proportions in scVelo ratios of splice to unspliced counts were calculated for each sample. It was done on filtered data after the removal of low-quality cells.

### RNA velocity

Based on the ratio of spliced versus unspliced counts across all genes in a cell, it is possible to predict the future transcriptional state of a given cell. Together, those predictions form a vector field of RNA velocity. Those velocity vectors are projected onto previously calculated embedding and averaged over the local neighbourhood of cells to provide the basis for the RNA velocity plots. The calculations are done with the scVelo package using the default stochastic model. Additionally, the basic velocity estimation was corrected using information from differential kinetics test using scVelo’s differential_kinetic_test function. This improves reliability of the predictions as our datasets represent a system with multiple lineages, where genes are likely to show different kinetic programmes across populations. Default parameters were used for all the calculations. velocity_embedding_stream function from scVelo was used for plotting on the embedding.

### GO term enrichment

DEGs with logFC>0.5 and FDR<0.05 were used for the analysis. GO term enrichment analysis was done with the goana function from the R package limma (Ritchie et al., 2015). The genes detected in a dataset (before filtering) were used as a background universe. The run was made with all default parameters. The p-values reported by goana are not adjusted for multiple testing, so we report as significant only GO terms with p-values greater than 10^−5^. For the plot’s terms were only selected if they had more than 50, and less than 2500 elements.

### Merging multiple datasets

The three datasets, healthy, healthy margin and ulcerated were combined for the purpose of joined analysis. QC and cell filtering were done separately for each sample in each dataset and only then data was concatenated and integrated together using the mutual nearest neighbours batch correction, just like for individual datasets. All other procedures were done analogically like for the healthy dataset.

### Cell lineage proportions

Cell lineage proportions (Figure 3C) were calculated per sample, as a percentage of cells in each lineage as compared to all cells from that sample. The stem, IA colonocytes and IA goblet lineages included only respective cell types. The Colonocytes, BEST4+ and Goblet lineages also included the TA and CT populations as separate data points. The height of a bar plot indicates the mean and whiskers show the interval of ± 1 standard error. Comparisons between groups are done with two sided Mann–Whitney *U* test. Only p-values smaller than 0.05 are reported.

### Lineage Venn diagram

To check for common DEGs a Venn diagram was created for the two populations, IA colonocytes and IA goblet in the ulcerated dataset. DEGs were calculated as compared to the rest of the cells in a lineage. Stem cells, TA colonocytes, colonocytes, CT Colonocytes and IA Colonocytes were extracted into a separate dataset. Then, DE analysis was performed as described before and genes with log2FC>1 and FDR<0.05 were used for the diagram. Analogous procedure was performed for the goblet lineage (including TA SOX4+ cells).

### Enrichment of transcription factor activity

To calculate activity of transcription factors we have used DoRothEA (Garcia-Alonso et al., 2019), a gene set resource containing signed transcription factors (known as regulons), and VIPER (Alvarez et al., 2016), a statistical method for inferencing protein activity by enriched regulon analysis. DoRothEA has been implemented in Python with a possibility of applying it to single-cell RNA-seq data (Holland et al., 2020). DoRothEA was run on log normalised expression using the dorothea.run function with the 3 most confident regulon sets (‘A’, ‘B’ and ‘C’) and all the default parameters.

### Pseudo-bulk pathway analysis

We have inferred pathway activities using the decoupler package and the PROGENy database (Badia-i-Mompel et al., 2022; Schubert et al., 2018). Analysis has been done according to the official vignette for pseudo-bulk functional analysis. In short, pseudo-bulk profiles have been generated from annotated raw count data. Each cell population in each sample was summarised into a single profile. Afterwards, data was normalised, log transformed, and then two conditions were contrasted using a t-test. Log fold changes between cell types were fed to the PROGENy model to estimate pathway activities with the consensus method.

### Molecular cartography

Biopsies from healthy individuals were fresh frozen in optimal cutting temperature (OCT compound, CellPath). Sections were stored at –80°C before use. For the analyses 10μm sections of the tissue were cut and placed onto a poly-lysine coated slide provided by Resolve Biosciences. Tissue was maintained frozen throughout for optimal RNA integrity. The slides were then shipped for analyses to Resolve Biosciences and processed on site. Thirty-one genes of interest were selected for design of transcript-specific probes. The genes were chosen based on differential expression analysis to abundantly mark the proliferative, differentiated and crypt top cells of the secretory lineage.

### Spatial transcriptomics visualisation

Spatial transcriptomics data from the Resolve Biosciences platform has been visualised using a proprietary ImageJ plugin Polylux provided by the company. Gene probes localisation was overlayed on top of a DAPI image of the tissue.

### Cell segmentation of spatial transcriptomics data

Cells in the spatial transcriptomics dataset have been segmented with a combination of two methods, image based cellpose and gene probes localisation based Baysor (Pachitariu and Stringer, 2022; Petukhov et al., 2022). The DAPI image of cell nuclei is segmented with cellpose, by first automatically detecting average cell diameter using the build-in ‘calibrate’ function. Afterwards, the cyto model is run with parameters flow set to 1 and cellprob set to –1.5. The resulting mask is saved as an image. Then, probe localisation is used to further improve the segmentation using Baysor. The tool is run on the x, y, and z coordinates of the gene probes, inputting the previously obtained mask as a prior to the model. Additionally, we specify the following parameters: i set to 1000, n-clusters set to 6, prior-segmentation-confidence set to 1 and force-2d. The resulting segmentation, together with gene probe localisation, is overlaid over the DAPI image for the final picture.

### Subdividing healthy margin dataset

To better discriminate cell states in the healthy margin, the dataset was subsetted based on CD74 expression. The threshold was set to 0.3970, which is the third quartile expression of CD74 in the healthy dataset. After subsetting the cells with log normalised counts, the processing pipeline, with batch correction, regression, dimensionality reduction, clustering and RNA velocity was repeated.

### Correlation matrix

The dendrogram and correlation matrix for the healthy, ulcerated and subdivided healthy margin datasets were calculated in scanpy using Pearson correlation on the first 50 principal components. Using a full feature matrix as well as other methods of correlation (Spearman, Kendall) gave similar results.

### Experimental acute colitis model

Acute colitis was induced in animals via the administration of dextran sodium sulfate (2.8% (w/v)) (DSS, MP Biomedicals; MW; 36.000-50.000) in the drinking water for 5 consecutive days *ad libitum* and the mice were subsequently transferred to normal drinking water. Control mice were provided normal drinking water. The severity of colitis during the acute phase was estimated by daily weight loss measurements, rectal bleeding, and stool consistency. Following euthanasia by cervical dislocation, colons were dissected, measured, and processed for the downstream analysis. Tissue was isolated and processed using standard techniques (Yui et al., 2018).

### Pharmacological inhibition of JAK-STAT in experimental acute colitis model

Tofacitinib was purchased from MedChemExpress (Cat. No.: HY-40354). The inhibitor was reconstituted in 10% DMSO (Sigma Aldrich), 40% PEG300 (MedChemExpress), 5% Tween-80 (Selleckchem) and 45% saline solution. Female C57BL/6J mice were divided into four experimental groups: group I (N=4) represented the healthy control that received normal drinking water from the beginning (D0) until the end of the experiment (D10), and a vehicle control from D5 – D10; group II (N=5) received normal drinking water throughout the course of the experiment, and were treated orally twice a day with tofacitinib (30 mg/kg) from D5 – D10; in group III (N=5; 1 mouse euthanised on D5 due to the severity of the disease) upon induction of experimental colitis (2,6% DSS (w/v)), on day 5 of the experiment mice were treated orally twice a day with vehicle and in group III (N=5; 1 mouse euthanised on D5 due to the severity of the disease) with tofacitinib (30 mg/kg). During the experiment, mice were weighed daily and on day 10 mice were euthanised by cervical dislocation and the tissue was collected for downstream analysis.

### Immunohistochemistry and fluorescence

For paraffin embedded tissues, sections were deparaffinized and antigen retrieval was performed by boiling in either freshly made sodium citrate (10 mM, pH 6) or Diva Decloaker 1X solution (Biocare Medical). Frozen samples were preserved in optimal cutting temperature (OCT compound, CellPath) and were cut (CM3050 Leica Microsystem), air-dried onto glass slides and stored at –80°C until immunofluorescence staining.

For antibody staining, sections were preincubated in blocking buffer containing BSA (1%) supplemented with adult bovine serum (10%), fish scale gelatin (0.5%) and TritonX100 (0.1%). Primary antibodies were incubated overnight at 4°C in blocking buffer. For immunohistochemistry, HRP-polymer secondary antibody incubation was performed at room temperature for 30-60 minutes in blocking buffer before DAB and hematoxylin counterstaining. For immunofluorescence, fluorophore-conjugated secondary antibodies were incubated at room temperature for 30-60 minutes in blocking buffer together with DAPI (Sigma-Adrich,1μg/ml). All washes were performed in Tween/PBS (0.2%). The following primary antibodies were used: anti-keratin 20 (KS.20.8, mouse monoclonal anti-human; Agilient-Dako), anti-PCNA (PC-10; Santa Cruz Biotechnology; #sc-56), anti-MUC2 (rabbit polyclonal anti-human, Santa Cruz Biotechnology; sc-15334), anti-CD74 (rabbit polyclonal; Atlas Antibodies HPA010592), anti-OLFM4 (rabbit polyclonal; Atlas Antibodies HPA077718), anti-CEACAM1 (rabbit polyclonal; Atlas Antibodies HPA011041), anti-TFF3 (rabbit polyclonal; Atlas Antibodies HPA035464), anti-β-catenin (mouse monoclonal, clone 14; BD Transduction Laboratories), anti-KI67 (MIB-1, mouse monoclonal; Agilient-Dako), anti-E-cadherin (mouse monoclonal, clone 36; BD Biosciences), anti-Reg1a (monoclonal rat antibody, clone 431211; R&D Systems), and secondary antibodies: Alexa Fluor (AF) 568 polyclonal donkey anti-mouse, Alexa Fluor (AF) 647 polyclonal donkey anti-rabbit, Alexa Fluor (AF) 488 polyclonal donkey anti-rat, AF555 polyclonal goat anti-mouse IgG1 antibody, AF647 polyclonal goat anti-mouse IgG2a antibody, AF488 donkey anti Rabbit IgG and AF647 polyclonal donkey anti-rabbit IgG antibody (all from Invitrogen) were used.

### Microscopy

Images of samples stained by immunohistochemistry was acquired using a NanoZoomer-XR Digital slide scanner (Hamamatsu C12000-01) and analysed in NDPview2 software. Images of sections stained by immunofluorescence were taken using laser scanning confocal microscope (Leica TSC SP8 or Leica Stellaris). Images were subsequently analysed in ImageJ, and Adobe Photoshop. Cell culture images was acquired using EVOS™ 7000 Imaging System (Invitrogen).

### Immunostaining of whole mount tissue

For whole mount staining PFA (Sigma) fixed tissue were cleared in CUBIC solution(Susaki et al., 2015) with DAPI (Sigma-Aldrich) for a week at 37°C. Before staining, the tissue was washed with PBS three times and then incubated with primary antibodies in BSA (1%) and TritonX100 (0.5%) in PBS at 4°C for three days. The tissue was washed 6 times in PBS, before incubated with secondary antibodies including DAPI (Sigma-Aldrich) in BSA (1%) and TritonX100 (0.5%) in PBS for three days at 4°C. After antibody staining the tissue was washed 6 times in PBS and before imaging the tissue was incubated in RapiClear (SUNJin Lab) for 30-60 minutes at room temperature. Whole mount images were taken using laser scanning confocal microscope (Leica TSC SP8). The following primary antibodies were used: anti-histone H3 (Ser10) (rabbit polyclonal; Cell Signal #9701) and anti-E-cadherin (mouse monoclonal, clone 36; BD Biosciences). The following secondary antibodies were use: AF647 polyclonal goat anti-mouse IgG2a antibody and AF488 Donkey anti Rabbit IgG (Invitrogen).

### Human organoids

Human colonic epithelial organoids were cultured using defined and optimized conditions for human colonic epithelium (Fujii et al., 2018). In this study, KRT20-tdTomato organoids, derived from healthy individuals and genetically modified to report KRT20 expression (CK20-IRES-iCaspase9-T2A-tdTomato-P2A-ERT2CreERT2) (Shimokawa et al., 2017; Sugimoto et al., 2018), were used to assess whether differentiated cells retain the capacity to self-renew and generate organoids. Organoids were prepared for sorting by dissociation into single cells with TrypLE (Invitrogen) incubation. tdTomato +/− cells were sorted on an ARIAIII (BD) instrument and RNA was isolated from sorted cells using RNeasy Micro Kit (Qiagen). Seeding efficiency was quantified by counting the number of structures forming after seeding as single cells.

### Cytokine and inhibitor treatment of human colon organoids

Human colonic organoids were treated with recombinant human IL-22 protein (R&D Systems) at 10 ng/mL, JAK inhibitor tofacitinib (MedChemExpress) at 50 μM and/or mTor inhibitor Rapamycin (Calbiochem) at 100 nM in normal organoid media without WNT and R-Spondin starting at day 4 after seeding as single cells, and RNA was isolated at day 7 using RNeasy Micro Kit (Qiagen). For each experiment untreated vehicle controls were applied; PBS for IL-22; DMSO for tofacitinib and Rapamycin. cDNA was generated using SuperScript III Reverse Transcriptase (Invitrogen) and Random primers (Thermo Fisher). RealQ Plus 2X Master Mix (Amplicon) was used for qPCR amplification on a QuantStudio 6 Flex (Applied Biosystems). For Western Blot, organoids were treated with IL-22 (10 ng/mL) and/or Rapamycin (100 nM) as described above including untreated control, and organoids were resuspended in RIPA protein lysis buffer containing protease inhibitor cocktail (Roche) at day 7. Thirty μg of protein extracts was separated on 4–12% Bis-Tris gels (Thermo Fisher) in MOPS running buffer and transferred to nitrocellulose membranes using the Mini Trans-Blot Cell system (Biorad). The membrane was blocked with 10% milk in Tris-buffered saline-Tween (TBS-T) solution one hour at room temperature and subsequently incubated overnight 4°C with primary antibodies: anti-S6 (1:10,000; 5G10 monoclonal rabbit; Cell Signaling 2217), anti-Phospho-S6 (1:10,000; D57.2.2E monoclonal rabbit; Cell Signaling 4858) and anti-a-Tubulin (1:10,000; 11H10 monoclonal rabbit; Cell Signaling) diluted in 10% milk TBS-T. Blots were washed three times in TBS-T followed by incubation with horseradish peroxidase (HRP)-conjugated secondary antibodies (Merck). Following washing in TBS-T, HRP-mediated substrate development was performed using Amersham detection system (ECL Western Blot detection kit, Merck). Blots were imaged using ChemiDoc™ MP Imaging System (Bio-Rad).

KRT20-tdTomato human colon organoids were treated with recombinant human IL-22 protein (R&D Systems) from day 4 to day 7 of organoid cultures in normal organoid media without WNT and R-Spondin. On day 7, cells were collected and resuspended in TrypLE to dissociate in single cells and flow sorted for differentiated (tdTomato+) and undifferentiated (tdTomato-) populations. Flow cytometry purified cells were seeded in Matrigel in normal human colon organoid media and at day 10 counted to assess the organoid forming efficiency.

### Whole-mount immunofluorescence of human colon organoids

The organoids were grown in Greiner CELLSTAR^®^ 96 well plates (Sigma-Aldrich), washed three times with 200 μl of PBS and centrifuged at 1811 *g* in a pre-cooled centrifuge at 10°C prior to fixation in 100 μL 4% PFA (Sigma) for 45 minutes at room temperature. After fixation organoids were washed with PBS, permeabilized with 100 μL of permeabilization buffer (0.5% TritonX-100 in PBS) for 1 hour with shaking at room temperature and counterstained with DAPI (Sigma-Aldrich, 10 μg/ml) in 0.1% TritonX-100 in PBS for 45 minutes. Following three washes in PBS, organoids were cleared with 50 μl of RapiClear 1.52 (SuNJin lab) for at least one hour before imaging.

### scRNA-seq of IL-22 treated human organoids

Human colonic organoids were derived from three healthy donors and cultured under standard condition in Matrigel and normal organoid media. Organoids were seeded as single cells and IL-22 treatment at 10 ng/mL was started on day 4. Treated and untreated (PBS) organoids were collected at day 7 and dissociated to single cells using TrypLE (Invitrogen) incubation at 37°C. Six different TotalSeq™ hashtag antibodies (A0251-A0256, Biolegend) were used to tag the individual samples (three untreated samples and three IL-22 treated samples). Each sample was incubated with 0.5 μg of a unique hashtag antibody for 20 minutes on ice. Before sorting the cells were incubated with DAPI for 10 minutes on ice. On an ARIAII FACS sorter (BD) DAPI negative cells were gated, and 6100 events/cells were sorted from each sample into the same tube for scRNA-sequencing. Single cell libraries were prepared using the 10X Genomics protocol v3.1 with implementation of hashtag libraries. To increase yield of hashtag-barcodes, an additive primer (HTO primer) was added in the step of cDNA amplification at 0.2 μM. 11 cycles were run for the cDNA amplification. After the cDNA amplification the hashtag-cDNA and endogenous cDNA is separated based on the size with SPRIselect beads (Beckman Coulter). After incubation with the SPRIselect beads, the hashtag-cDNA was in the supernatant containing smaller fragments (which is normally discarded) and was saved for the hashtag-library following the protocol from Stoeckius et al. (Stoeckius et al., 2018) and CiteSeq (https://cite-seq.com). In short, the supernatant was cleaned using SPRIselect beads and 30 ng of cDNA is used as input for the PCR amplification (20 cycles) with Kapa HiFi HotStart Ready Mix (Roche Diagnostics) and addition of Illumina-index sequences. To generate the library of the endogenous cDNA the 10X protocol was followed. The dual index kit set AA (10X genomics) were used for sample indexing and 11 cycles were used during the final amplification step. The hashtag cDNA and endogenous cDNA libraries were diluted to 4 nM and pooled (5% hashtag + 95 % endogenous cDNA) before being sequenced on a NextSeq2000 sequencer (Illumina).

### Analysis of organoids scRNA-seq data

scRNA-seq data from human colon organoids was analysed in the same manner as described before, apart from the following changes. Count files were not pre-processed for RNA velocity measurement using velocyto. Data was generated with the Cell Hashing technique, which uses oligo-tagged antibodies against surface proteins to barcode single cells. This allows for samples to be multiplexed together and run in a single experiment. The data was demultiplexed using the HTODemux() function from Seurat (Hao et al., 2021). As most doublets are removed during the demultiplexing process, we did not perform any additional doublet filtering. General mitochondrial content was lower, so cells with more than 0.15 mitochondrial features were filtered out. Differences between samples were batch corrected using the Harmony tool implemented into scanpy (Korsunsky et al., 2019). Neighbourhood graph was calculated using the default method in scanpy, instead of the bbknn method. Cell types were annotated using unsupervised clustering and gene markers found in the patient dataset.

### REG1A-mNeonGreen human organoid reporter

sgRNAs targeting the C-terminal end of REG1A were designed and cloned into pSPgRNA (Addgene #47108) according to the protocol from Ran et al. (Ran et al., 2013). For tagging the CRISPR-HOT method published from Artegiani et al. (Artegiani et al., 2020) was used. For electroporation the protocol published by Fujii et al. (Fujii et al., 2015) was used. In short, organoids were seeded in high density with normal organoid media. Two days before electroporation Rpso and Wnt were removed and Chir99021 (3 μM)) and rock inhibitor Y-27632 (10 μM) was added. One day before electroporation DMSO (1.25%) was added to the media. On the day of electroporation organoids were dissociated into small clumps of 3-5 cells using mechanical pipetting and TrypLE incubation and resuspended in Opti-MEM Reduced Serum Medium (Gibco). 5 μg of the cloned REG1A sgRNA-plasmid, 5 μg pCAS9-mCherry-Frame+1 (Addgene #66940), and 5 μg of pCRISPR-HOT_mNEON plasmid (a kind gift from Hans Clevers) were electroporated into the cells using the NEPA2000 electroporating system and hereafter seeded in Matrigel with normal organoid media supplemented with rock inhibitor Y-27632 (10 μM). Three days later, organoids were dissociated and mCherry positive cells (marking successful electroporation of the pCAS9-mCherry-Frame+1 plasmid) were sorted on ARIAIII sorter (BD Biosciences) and seeded in Matrigel with normal organoid media supplemented with rock inhibitor Y-27632 (10 μM). Organoids were passaged and seeded as single cells and on day 4 treated with IL-22 (10 ng/mL) for three days to induce REG1A-mNeonGreen expression. Single mNeonGreen+ organoids were picked and seeded in Matrigel to obtain clonal lines. DNA was harvested from clonal lines and extracted using DNeasy Blood & Tissue kit (Qiagen) and a PCR was run to amplify the C-terminal end of *REG1A*. DNA was extracted from the agarose gel using QIAquick Gel Extraction Kit (Qiagen) and send for Sanger sequencing to verify in-frame insertion of the mNeonGreen construct. Clones 3 and 4 had correct insertion and were used for further experiments.

### Seeding efficiency and qPCR of FACS sorted REG1A-mNeonGreen organoids

The REG1A-mNeonGreen organoids were seeded as single cells and treatment with IL-22 (10 ng/mL) was started on day 4. Organoids were dissociated to single cells on day 7 with mechanical pipetting and TrypLE incubation. mNeonGreen negative and positive cells were sorted on a FACSymphony™ S6 sorter (BD Biosciences). WT organoids untreated and treated with IL-22 in the same experiment was used as control for gating. 20,000 events/cells were sorted directly into RNA lysis buffer and RNA was extracted using RNeay Micro Kit (Qiagen). cDNA synthesis and qPCR were set up as described above. For organoid seeding efficiency, 16,000 events/cells were sorted into normal organoid media supplemented rock inhibitor Y-27632 (10 μM). The sorted cells were seeded in Matrigel in four wells in a 96-well plate and cultured in normal organoid media supplemented rock inhibitor Y-27632 (10 μM). Z-stack images were taking on day 6-7 after seeding using EVOS™ 7000 Imaging System (Invitrogen). and used for quantification of organoids.

### Statistical Methods

The number of biological and technical replicates and the number of animals is indicated in figure legends. All tested animals were included. Sample size was not predetermined. Experiments were performed without methods of randomization or blinding. For all experiments with error bars, the standard error of the mean (SEM) was calculated to indicate the variation within each experiment. Statistics analysis was performed in Prism and R. Non-parametric t-tests were used for the comparison between two different conditions. For experiments with more than two conditions, ANOVA test was used to calculate significance. Annotation for p values in figure legends regardless of statistical test type are: ∗p < 0.05, ∗∗p < 0.01, ∗∗∗p < 0.001, ∗∗∗∗p < 0.0001.

## ACKNOWLEDGMENTS

We thank Hans Clevers, Calvin Kuo and Chris Garcia for kindly sharing cell culture reagents and plasmids; members of the Jensen, Nielsen labs and Viktor Petukhov for comments and suggestions; the reNEW Platforms and BRIC imaging and histology core for technical expertise and support. We further thank staff at the Endoscopy Unit, Department of Gastroenterology and Department of Pathology, Herlev Hospital for assistance with handling of patient tissue samples. The graphical abstract was created using BioRender. KK and AMB are recipients of a fellowship from the Copenhagen Bioscience Ph.D. Programme (NNF18CC0033666) and Danish Diabetes Academy (NNF17SA0031406). This work was supported by the Independent Research Fund Denmark (0134-00111B to KBJ), the Novo Nordisk Foundation (NNF17OC0028730; NNF18OC0034066; NNF20OC0064376 to KBJ), the Leo Pharma Foundation (LF-OC-19-000169) and the European Union’s Horizon 2020 research and innovation programme (STEMHEALTH ERCCoG682665 to KBJ). The Novo Nordisk Foundation Center for Stem Cell Medicine is supported by the Novo Nordisk Foundation (NNF21CC0073729).

## AUTHOR CONTRIBUTIONS

OHN and KBJ conceived the project. GM, SLH, KK, LK, SU, MM, AMB, TS, OHN and KBJ designed experiments. LK, LBR, and OHN collected and analyzed clinical samples. GM, SLH, KK, LK, SU, MM, AMB performed experiments. GM analyzed scSEQ data. KKH assisted with scSEQ analysis. KBJ wrote the manuscript with input from GM, SLH, KK, LK and OHN; Funding Acquisition, OHN and KBJ; Supervision, OHN and KBJ.

## Supplementary materials

**Supplementary Figure 1.**
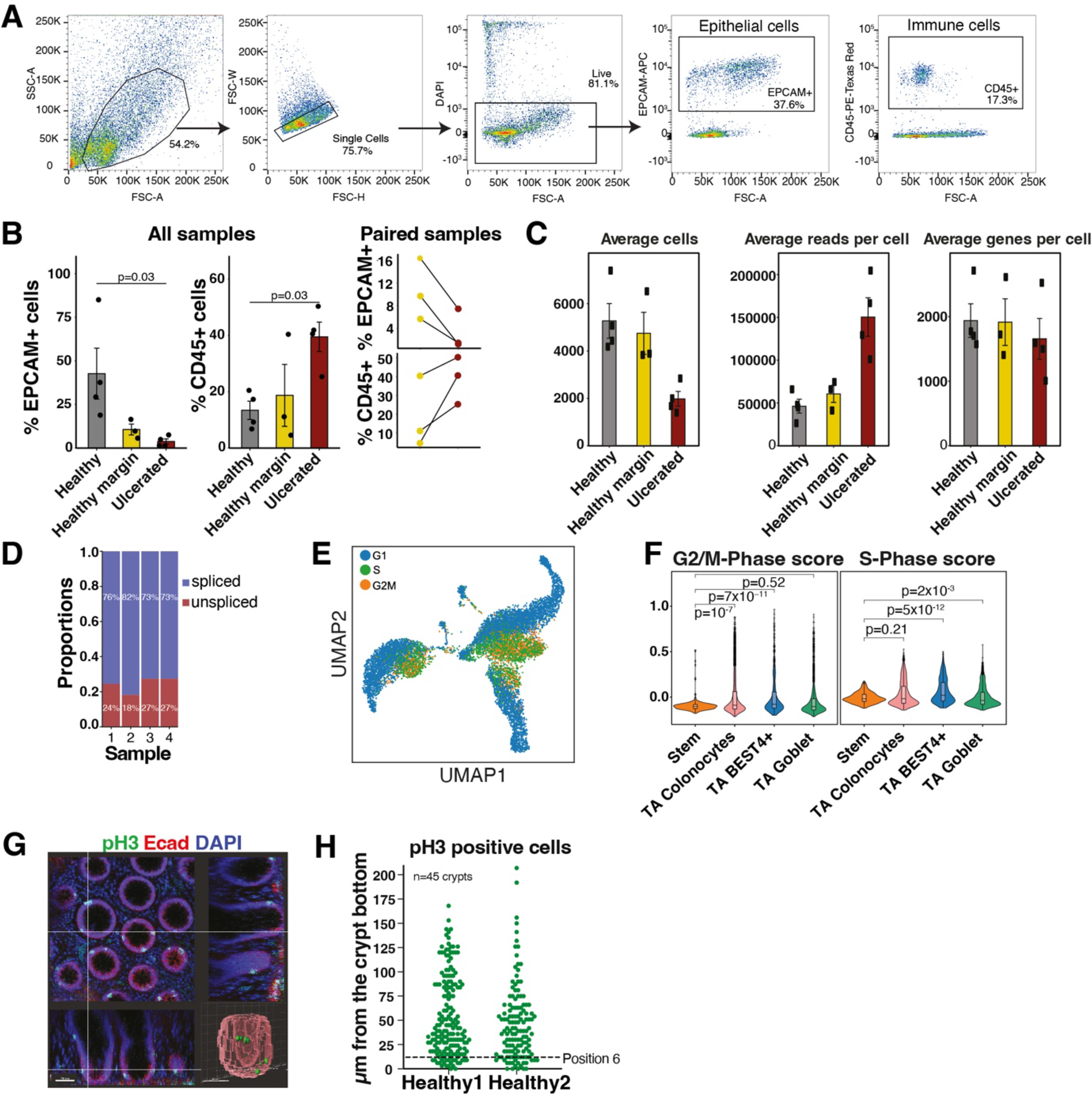
Single cell profiling of healthy colonic epithelium. (*A*) Flow cytometry sorting strategy for isolation of epithelial cells from clinical samples based on markers for live-dead, EPCAM+ and CD45+. (*B*) Average fraction of EPCAM and CD45 positive cells in the clinical samples analysed (left) with a paired analysis of biopsies from the UC patients from ulcerated and healthy margin regions (right). Error bars indicates the S.E.M. and p-value tested with one-sided Mann–Whitney U test is indicated. (*C*) Quality control statistics of samples after 10x Cell Ranger processing of the separate runs. Error bars indicates the S.E.M. (*D*) Proportions of spliced/unspliced counts among the healthy samples. (*E*) UMAP plot of cells in the healthy dataset colored by cell cycle. (*F*) Violin plot of individual cell scores for the cell cycle phase signatures in selected cell populations. Significance tested with one-sided Mann–Whitney U test. (*G*) Immunostaining of whole mount healthy human tissue for phospho-Histone-3 (pH3) (green) and E-Cadherin (red). Scale bar: 50 µm. Insert: 3D reconstruction of a colonic crypt. Scale bar: 40 µm. (*H*) Quantification of the distance (µm) from the bottom of colonic crypts to pH3 positive cells in two healthy human colonic samples. Cell position 6 is marked with a dotted line. N = 45 crypts quantified in total.

**Supplementary Figure 2.**
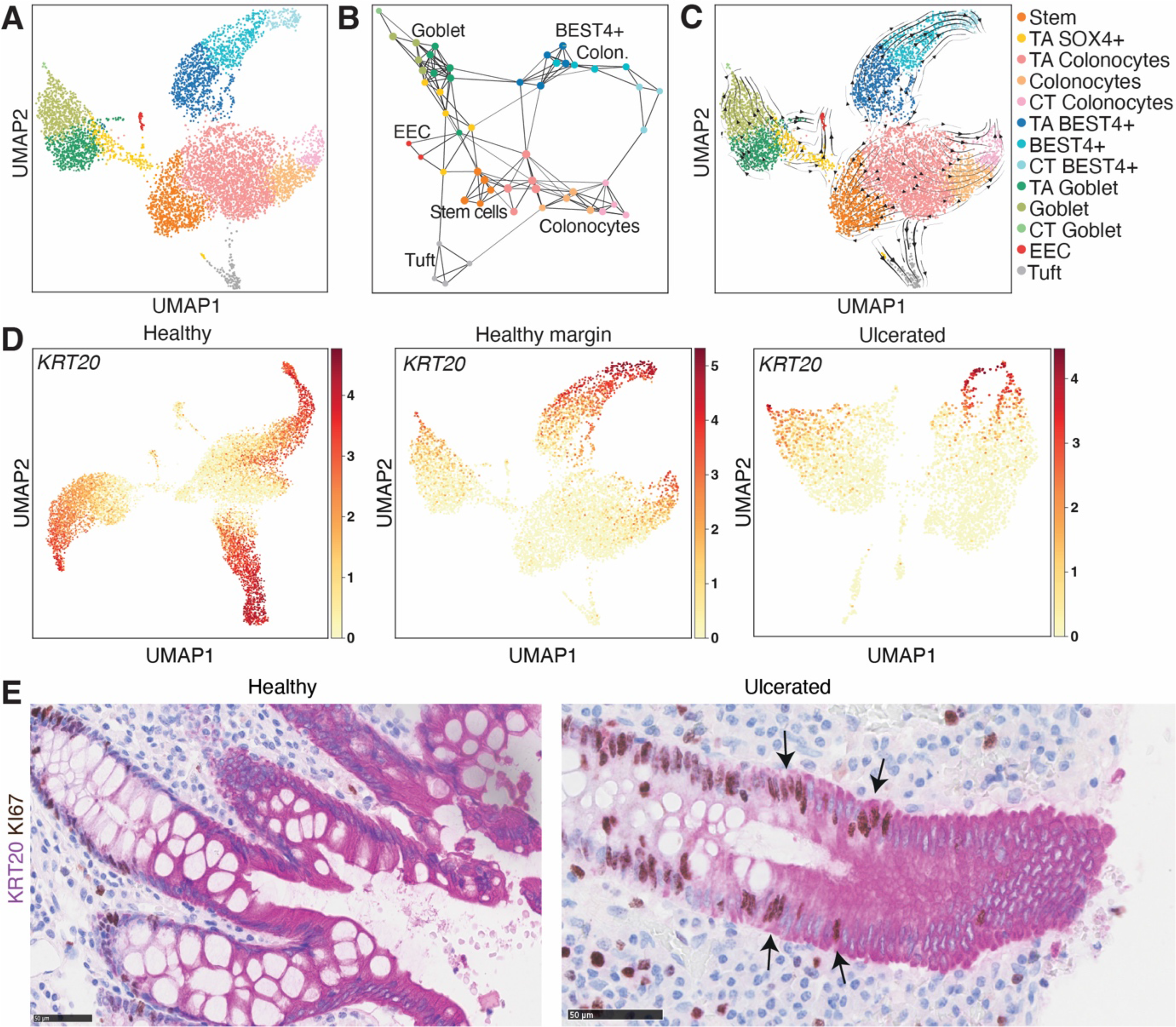
Single cell profiling of healthy margin of colonic epithelium from UC patients. (*A*) UMAP of 6,762 colonic epithelial cells from the healthy margin of UC patients annotated based on marker expression. (*B*) Cell type trajectories based on over-clustering. Nodes represent clusters colored by cell type of origin, while edges indicate high degree of similarities. (*C*) RNA velocity measurements projected onto the UMAP embedding shown in (A). (*D*) UMAP plots with heatmaps of normalized expression of the *KRT20* gene in healthy, healthy margin and ulcerated data sets. *(E)* Higher magnification of representative IHC for KRT20 and KI67 in healthy and ulcerated human colon. Arrows indicate KRT20 and Ki67 double positive cells. Scale bar: 50 µm.

**Supplementary Figure 3.**
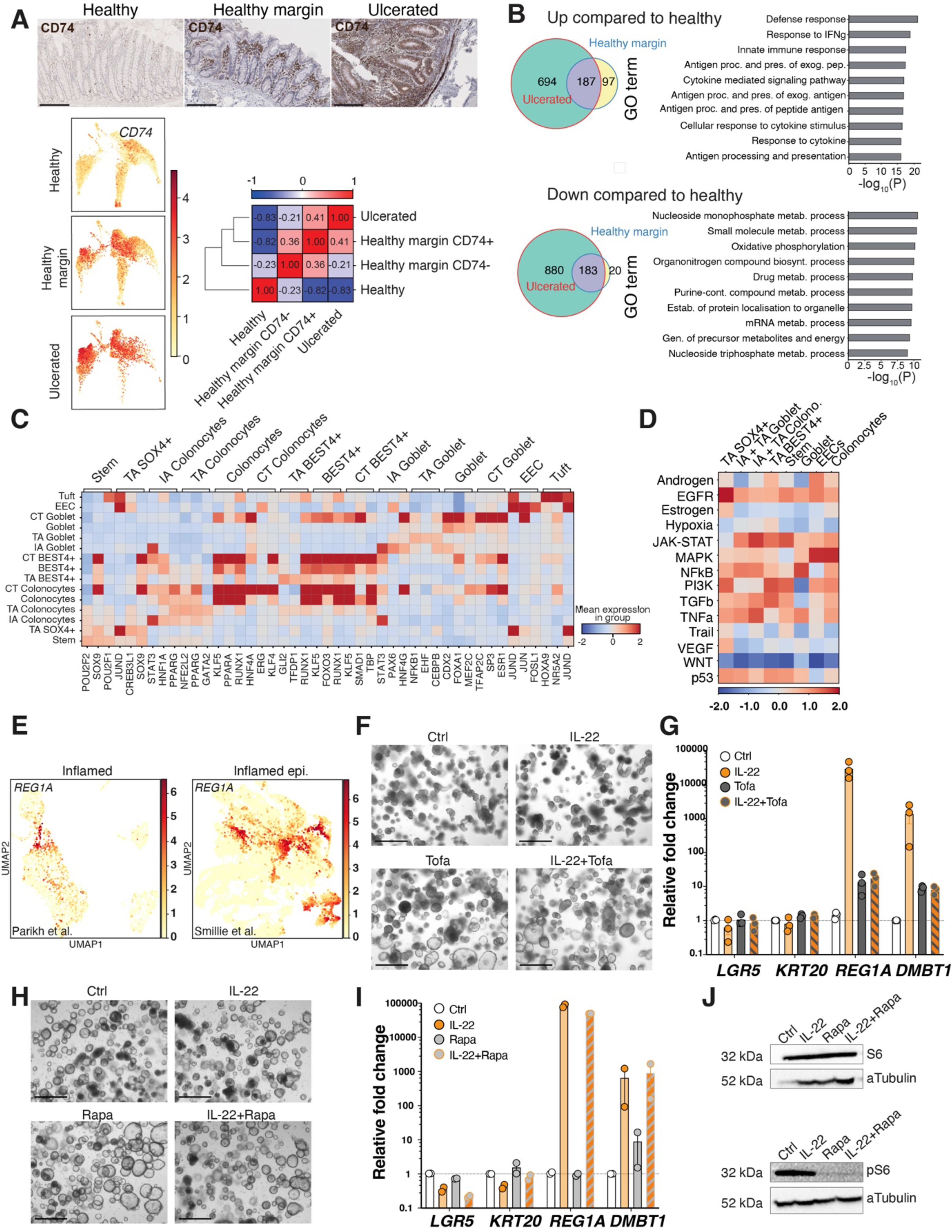
The IA cell states are induced through STAT signalling. *(A)* Immunostaining for CD74 in healthy, healthy-margin and ulcerated tissues. Scale bars: 100 µm. UMAP plots with heatmaps of normalized expression of the CD74 gene split into 3 panels based on a dataset of origin. Similarity correlation matrix between healthy, ulcerated and healthy-margin (split on the CD74 expression with cut-off value for the split based on the third quartile of CD74 expression in the healthy dataset) samples calculated on first 50 principal components using Pearson correlation showing that there is high correlation between the CD74+ cells from the non-ulcerated samples and the ulcerated state. This is also supported by the dendrogram visualizing hierarchical clustering on the groups. *(B)* Venn diagrams of the overlap between genes upregulated and downregulated by stem cells in the ulcerated and healthy-margin (when compared to the healthy state). This reveals a much larger than expected overlap in fraction of shared genes. Top 10 terms from GO term overrepresentation analysis of the shared differentially expressed genes. (*C*) Heatmap of scaled transcription factor activity of DoRothEA regulons. Top three differentially active regulons for each of the cell type clusters are shown. (*D*) Heatmap of scaled pathway activity for ulcerated cell populations. Activities have been calculated using genes differentially expressed in the ulcerated dataset, as compared to the healthy dataset. The DEGs have been found between pseudo-bulk profiles of cell types from the two datasets. Pseudo-bulk profiles have been generated only if at least two biological replicates per condition were available. In the ulcerated dataset, IA Colonocytes and IA Goblet cells have been combined with TA Colonocytes and TA Goblet cells, respectively. (*E*) UMAP plots with heatmaps of normalized expression of the *REG1A* gene in inflamed epithelial data sets from Parikh et al. 2019 and Smillie et al. 2019 studies. (*F*) Representative images of WT human colonic organoids control treated or treated with IL-22 and/or tofacitinib (Tofa) for 3 days. Scale bars = 650 μm. (*G*) Expression levels of indicates genes of WT organoids control treated or treated with IL-22 and/or tofacitinib (Tofa) for 3 days normalized to *GAPDH* expression. Fold change relative to control treated organoids ± SEM; n=3. (*H*) Representative images of WT human colonic organoids control treated or treated with IL-22 and/or rapamycin (Rapa) for 3 days. Scale bars = 650 μm. (*I*) Expression levels of indicates genes of WT organoids control treated or treated with IL-22 and/or rapamycin (Rapa) for 3 days normalized to *GAPDH* expression. Fold change relative to control treated organoids ± SEM; n=2. (*J*) Western blot showing protein levels of S6 (top) and phosphorylated S6 (bottom) in WT human colonic organoids control treated or treated with IL-22 and/or rapamycin (Rapa) for 3 days. α-Tubulin was used as a loading control.

**Supplementary Figure 4.**
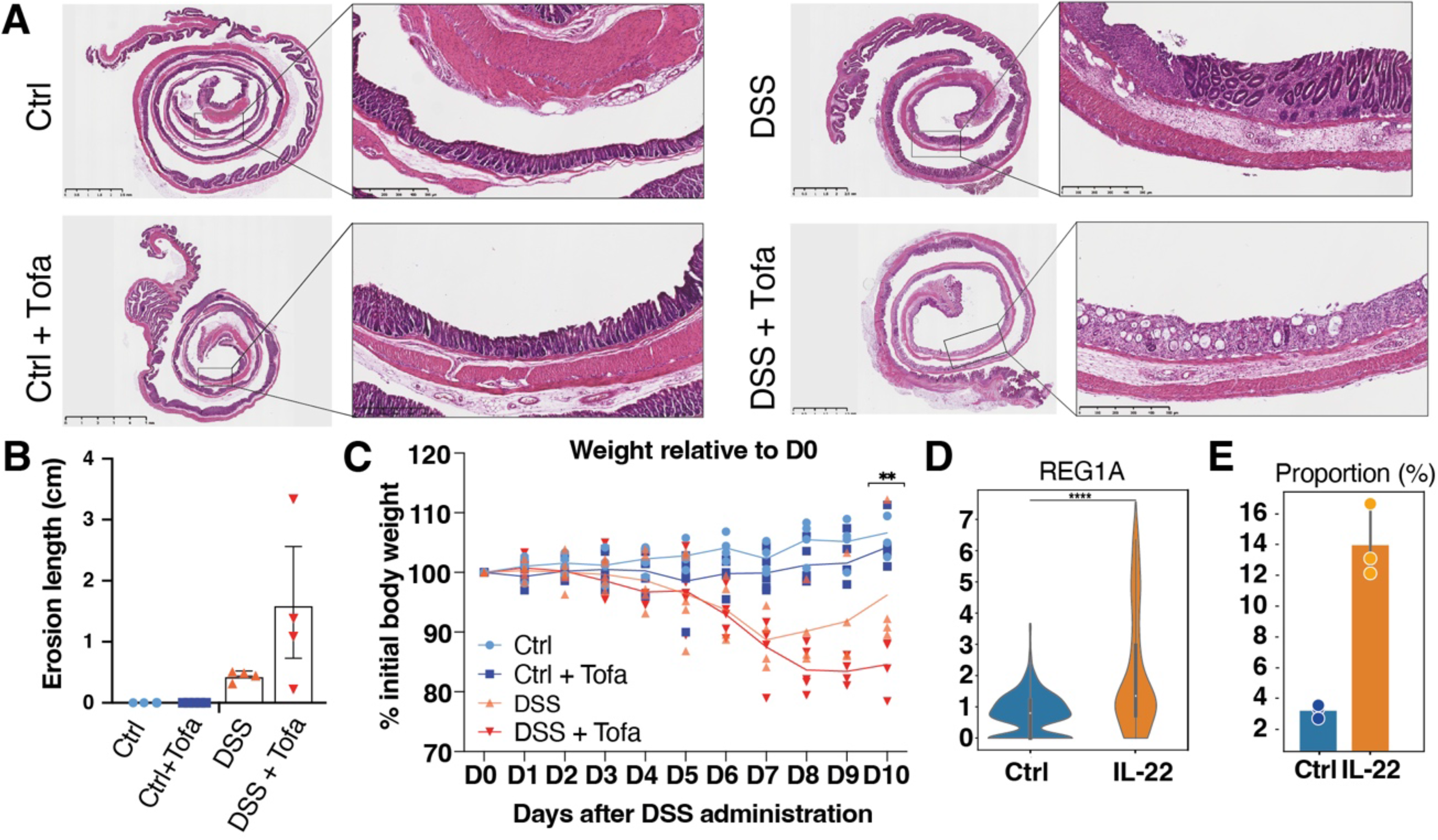
Characterisation of the role of JAK/STAT signalling in tissue regeneration and *REG1* expression. (*A*) H&E staining of sections of Swiss rolls of the entire mouse colon of control and DSS treated mice with tofacitinib (Tofa) or vehicle treatment from D5-D10. (*B*) Histological assessment of the length of epithelial erosions from H&E stainings *(A)*. For CTRL, N=3; for CTRL + tofacitinib. N=5; for DSS, N=4; for DSS + tofacitinib, N=4. *(C)* Body weight changes of mice in all experimental groups. Mice were treated with DSS from D0-D5 and Tofacitinib or vehicle control from D5-D10. (D) Violin plot with normalized *REG1A* gene expression in cells in the control and IL-22 treated organoid datasets. The boxplot shows the quartiles of the dataset while the whiskers extend to show the rest of the distribution, except for outliers. Significance tested with two-sided Mann–Whitney U test. (*E*) Bar plot of percentage of *REG1A* positive cells (normalized expression in the third quantile) in samples in the control and IL-22 treated organoid datasets. Error bars show 95% confidence interval.

**Supplementary Figure 5.**
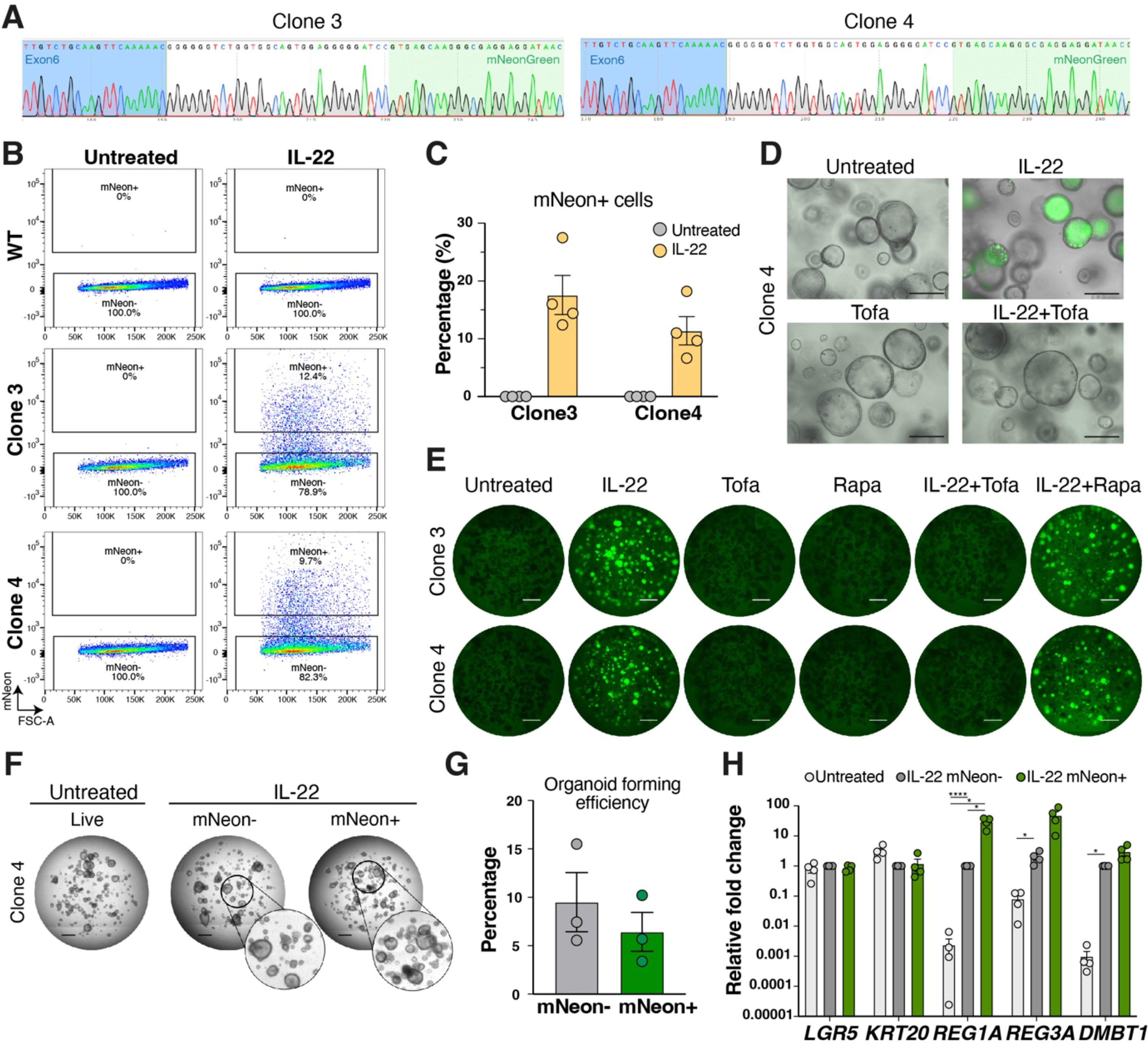
*REG1A* expressing cells exhibit regenerative capabilities. (*A*) Sanger sequencing at the site of insertion of the mNeonGreen tag at the C-terminal of *REG1A* in Clone 3 and Clone 4. (*B*) Representative flow cytometry analysis of mNeonGreen in WT and REG1A-mNeonGreen Clone 3 and Clone 4 in untreated and IL-22 treated conditions. (*C*) Quantification of mNeonGreen positive cells in REG1A-mNeonGreen Clone 3 and Clone 4 cells untreated or treated with IL-22 for three days. Data shown as ± SEM and n=4. (*D*) REG1A-mNeonGreen Clone 4 untreated or treated with IL-22 and/or tofacitinib (Tofa) for three days. Scale bars: 275 µm. (*E*) Representative images of REG1A-mNeonGreen Clone 3 and 4 untreated or treated with IL-22, and/or tofacitinib (Tofa), and/or rapamycin (Rapa) for three days. Scale bars: 1mm. (*F*) Representative images of REG1A-mNeonGreen Clone 4 organoids re-seeded in Matrigel after FACS sorting, either untreated (live cells) or treated with IL-22 for three days (mNeonGreen+ and mNeonGreen-cells). Scale bars: 1mm. (*G*) Quantification of organoid forming efficiency related to F. Data shown as ± SEM; n=3 for each population with average of four wells counted at each experiment. (*H*) Expression levels of indicates genes of FACS sorted cells from REG1A-mNeonGreen (clone 4) untreated control cells (light grey) and IL-22 treated cells, either mNeonGreen-(dark grey) or mNeonGreen+ cells (green), normalized to *GAPDH* expression. Fold change relative to mNeonGreen-cells ± SEM; n=4 for each population. *p< 0.05, **p < 0.01, ***p < 0.001, ****p < 0.0001; two-way ANOVA with Tukey’s multiple comparison test.

## REFERENCES

1. Alvarez, M.J., Shen, Y., Giorgi, F.M., Lachmann, A., Ding, B.B., Ye, B.H., and Califano, A. (2016). Functional characterization of somatic mutations in cancer using network-based inference of protein activity. Nat Genet 48, 838–847. 10.1038/ng.3593.

2. Artegiani, B., Hendriks, D., Beumer, J., Kok, R., Zheng, X., Joore, I., Chuva de Sousa Lopes, S., van Zon, J., Tans, S., and Clevers, H. (2020). Fast and efficient generation of knock-in human organoids using homology-independent CRISPR-Cas9 precision genome editing. Nat Cell Biol 22, 321–331. 10.1038/s41556-020-0472-5.

3. Badia-i-Mompel, P., Vélez Santiago, J., Braunger, J., Geiss, C., Dimitrov, D., Müller-Dott, S., Taus, P., Dugourd, A., Holland, C.H., Ramirez Flores, R.O., and Saez-Rodriguez, J. (2022). decoupleR: ensemble of computational methods to infer biological activities from omics data. Bioinformatics Advances 2. 10.1093/bioadv/vbac016.

4. Bergen, V., Lange, M., Peidli, S., Wolf, F.A., and Theis, F.J. (2020). Generalizing RNA velocity to transient cell states through dynamical modeling. Nat Biotechnol 38, 1408–1414. 10.1038/s41587-020-0591-3.

5. Beumer, J., and Clevers, H. (2021). Cell fate specification and differentiation in the adult mammalian intestine. Nat Rev Mol Cell Biol 22, 39–53. 10.1038/s41580-020-0278-0.

6. Corridoni, D., Antanaviciute, A., Gupta, T., Fawkner-Corbett, D., Aulicino, A., Jagielowicz, M., Parikh, K., Repapi, E., Taylor, S., Ishikawa, D., et al. (2020). Single-cell atlas of colonic CD8(+) T cells in ulcerative colitis. Nat Med 26, 1480–1490. 10.1038/s41591-020-1003-4.

7. Cox, C.B., Storm, E.E., Kapoor, V.N., Chavarria-Smith, J., Lin, D.L., Wang, L., Li, Y., Kljavin, N., Ota, N., Bainbridge, T.W., et al. (2021). IL-1R1-dependent signaling coordinates epithelial regeneration in response to intestinal damage. Sci Immunol 6. 10.1126/sciimmunol.abe8856.

8. Danese, S., Vermeire, S., Zhou, W., Pangan, A.L., Siffledeen, J., Greenbloom, S., Hebuterne, X., D’Haens, G., Nakase, H., Panes, J., et al. (2022). Upadacitinib as induction and maintenance therapy for moderately to severely active ulcerative colitis: results from three phase 3, multicentre, double-blind, randomised trials. Lancet 399, 2113–2128. 10.1016/S0140-6736(22)00581-5.

9. de Sousa, E.M.F., and de Sauvage, F.J. (2019). Cellular Plasticity in Intestinal Homeostasis and Disease. Cell Stem Cell 24, 54–64. 10.1016/j.stem.2018.11.019.

10. Denk, H., Lackinger, E., Zatloukal, K., and Franke, W.W. (1987). Turnover of cytokeratin polypeptides in mouse hepatocytes. Exp Cell Res 173, 137–143. 10.1016/0014-4827(87)90339-9.

11. Elmentaite, R., Ross, A.D.B., Roberts, K., James, K.R., Ortmann, D., Gomes, T., Nayak, K., Tuck, L., Pritchard, S., Bayraktar, O.A., et al. (2020). Single-Cell Sequencing of Developing Human Gut Reveals Transcriptional Links to Childhood Crohn’s Disease. Dev Cell 55, 771–783 e775. 10.1016/j.devcel.2020.11.010.

12. Fawkner-Corbett, D., Antanaviciute, A., Parikh, K., Jagielowicz, M., Geros, A.S., Gupta, T., Ashley, N., Khamis, D., Fowler, D., Morrissey, E., et al. (2021). Spatiotemporal analysis of human intestinal development at single-cell resolution. Cell 184, 810–826 e823. 10.1016/j.cell.2020.12.016.

13. Fujii, M., Matano, M., Nanki, K., and Sato, T. (2015). Efficient genetic engineering of human intestinal organoids using electroporation. Nat Protoc 10, 1474–1485. 10.1038/nprot.2015.088.

14. Fujii, M., Matano, M., Toshimitsu, K., Takano, A., Mikami, Y., Nishikori, S., Sugimoto, S., and Sato, T. (2018). Human Intestinal Organoids Maintain Self-Renewal Capacity and Cellular Diversity in Niche-Inspired Culture Condition. Cell Stem Cell 23, 787–793 e786. 10.1016/j.stem.2018.11.016.

15. Garcia-Alonso, L., Holland, C.H., Ibrahim, M.M., Turei, D., and Saez-Rodriguez, J. (2019). Benchmark and integration of resources for the estimation of human transcription factor activities. Genome Res 29, 1363–1375. 10.1101/gr.240663.118.

16. Gehart, H., and Clevers, H. (2019). Tales from the crypt: new insights into intestinal stem cells. Nat Rev Gastroenterol Hepatol 16, 19–34. 10.1038/s41575-018-0081-y.

17. Graham, D.B., and Xavier, R.J. (2020). Pathway paradigms revealed from the genetics of inflammatory bowel disease. Nature 578, 527–539. 10.1038/s41586-020-2025-2.

18. Granau, A.M., Boye, T.L., Jensen, K.B., and Nielsen, O.H. (2023). Deucravacitinib (Sotyktu) for plaque psoriasis. Trends Pharmacol Sci 44, 252–253. 10.1016/j.tips.2023.01.004.

19. Gregorieff, A., Liu, Y., Inanlou, M.R., Khomchuk, Y., and Wrana, J.L. (2015). Yap-dependent reprogramming of Lgr5(+) stem cells drives intestinal regeneration and cancer. Nature 526, 715–718. 10.1038/nature15382.

20. Hao, Y., Hao, S., Andersen-Nissen, E., Mauck, W.M., 3rd, Zheng, S., Butler, A., Lee, M.J., Wilk, A.J., Darby, C., Zager, M., et al. (2021). Integrated analysis of multimodal single-cell data. Cell 184, 3573–3587 e3529. 10.1016/j.cell.2021.04.048.

21. Harnack, C., Berger, H., Antanaviciute, A., Vidal, R., Sauer, S., Simmons, A., Meyer, T.F., and Sigal, M. (2019). R-spondin 3 promotes stem cell recovery and epithelial regeneration in the colon. Nat Commun 10, 4368. 10.1038/s41467-019-12349-5.

22. He, G.W., Lin, L., DeMartino, J., Zheng, X., Staliarova, N., Dayton, T., Begthel, H., van de Wetering, W.J., Bodewes, E., van Zon, J., et al. (2022). Optimized human intestinal organoid model reveals interleukin-22-dependency of paneth cell formation. Cell Stem Cell 29, 1333–1345 e1336. 10.1016/j.stem.2022.08.002.

23. Hindryckx, P., Jairath, V., and D’Haens, G. (2016). Acute severe ulcerative colitis: from pathophysiology to clinical management. Nat Rev Gastroenterol Hepatol 13, 654–664. 10.1038/nrgastro.2016.116.

24. Holland, C.H., Tanevski, J., Perales-Paton, J., Gleixner, J., Kumar, M.P., Mereu, E., Joughin, B.A., Stegle, O., Lauffenburger, D.A., Heyn, H., et al. (2020). Robustness and applicability of transcription factor and pathway analysis tools on single-cell RNA-seq data. Genome Biol 21, 36. 10.1186/s13059-020-1949-z.

25. Ishikawa, K., Sugimoto, S., Oda, M., Fujii, M., Takahashi, S., Ohta, Y., Takano, A., Ishimaru, K., Matano, M., Yoshida, K., et al. (2022). Identification of Quiescent LGR5(+) Stem Cells in the Human Colon. Gastroenterology 163, 1391–1406 e1324. 10.1053/j.gastro.2022.07.081.

26. Kinchen, J., Chen, H.H., Parikh, K., Antanaviciute, A., Jagielowicz, M., Fawkner-Corbett, D., Ashley, N., Cubitt, L., Mellado-Gomez, E., Attar, M., et al. (2018). Structural Remodeling of the Human Colonic Mesenchyme in Inflammatory Bowel Disease. Cell 175, 372–386 e317. 10.1016/j.cell.2018.08.067.

27. Kobayashi, T., Siegmund, B., Le Berre, C., Wei, S.C., Ferrante, M., Shen, B., Bernstein, C.N., Danese, S., Peyrin-Biroulet, L., and Hibi, T. (2020). Ulcerative colitis. Nat Rev Dis Primers 6, 74. 10.1038/s41572-020-0205-x.

28. Korsunsky, I., Millard, N., Fan, J., Slowikowski, K., Zhang, F., Wei, K., Baglaenko, Y., Brenner, M., Loh, P.R., and Raychaudhuri, S. (2019). Fast, sensitive and accurate integration of single-cell data with Harmony. Nat Methods 16, 1289–1296. 10.1038/s41592-019-0619-0.

29. La Manno, G., Soldatov, R., Zeisel, A., Braun, E., Hochgerner, H., Petukhov, V., Lidschreiber, K., Kastriti, M.E., Lonnerberg, P., Furlan, A., et al. (2018). RNA velocity of single cells. Nature 560, 494–498. 10.1038/s41586-018-0414-6.

30. Larsen, H.L., and Jensen, K.B. (2021). Reprogramming cellular identity during intestinal regeneration. Curr Opin Genet Dev 70, 40–47. 10.1016/j.gde.2021.05.005.

31. Lee-Six, H., Olafsson, S., Ellis, P., Osborne, R.J., Sanders, M.A., Moore, L., Georgakopoulos, N., Torrente, F., Noorani, A., Goddard, M., et al. (2019). The landscape of somatic mutation in normal colorectal epithelial cells. Nature 574, 532–537. 10.1038/s41586-019-1672-7.

32. Lindemans, C.A., Calafiore, M., Mertelsmann, A.M., O’Connor, M.H., Dudakov, J.A., Jenq, R.R., Velardi, E., Young, L.F., Smith, O.M., Lawrence, G., et al. (2015). Interleukin-22 promotes intestinal-stem-cell-mediated epithelial regeneration. Nature 528, 560–564. 10.1038/nature16460.

33. McGinnis, C.S., Murrow, L.M., and Gartner, Z.J. (2019). DoubletFinder: Doublet Detection in Single-Cell RNA Sequencing Data Using Artificial Nearest Neighbors. Cell Syst 8, 329–337 e324. 10.1016/j.cels.2019.03.003.

34. Merlos-Suarez, A., Barriga, F.M., Jung, P., Iglesias, M., Cespedes, M.V., Rossell, D., Sevillano, M., Hernando-Momblona, X., da Silva-Diz, V., Munoz, P., et al. (2011). The intestinal stem cell signature identifies colorectal cancer stem cells and predicts disease relapse. Cell Stem Cell 8, 511–524. 10.1016/j.stem.2011.02.020.

35. Nicholson, A.M., Olpe, C., Hoyle, A., Thorsen, A.S., Rus, T., Colombe, M., Brunton-Sim, R., Kemp, R., Marks, K., Quirke, P., et al. (2018). Fixation and Spread of Somatic Mutations in Adult Human Colonic Epithelium. Cell Stem Cell 22, 909–918 e908. 10.1016/j.stem.2018.04.020.

36. Nielsen, O.H., Boye, T.L., Gubatan, J., Chakravarti, D., Jaquith, J.B., and LaCasse, E.C. (2023). Selective JAK1 inhibitors for the treatment of inflammatory bowel disease. Pharmacol Ther, 108402. 10.1016/j.pharmthera.2023.108402.

37. Nusse, Y.M., Savage, A.K., Marangoni, P., Rosendahl-Huber, A.K.M., Landman, T.A., de Sauvage, F.J., Locksley, R.M., and Klein, O.D. (2018). Parasitic helminths induce fetal-like reversion in the intestinal stem cell niche. Nature 559, 109–113. 10.1038/s41586-018-0257-1.

38. Oshima, H., Kok, S.Y., Nakayama, M., Murakami, K., Voon, D.C., Kimura, T., and Oshima, M. (2019). Stat3 is indispensable for damage-induced crypt regeneration but not for Wnt-driven intestinal tumorigenesis. FASEB J 33, 1873–1886. 10.1096/fj.201801176R.

39. Pachitariu, M., and Stringer, C. (2022). Cellpose 2.0: how to train your own model. Nat Methods 19, 1634–1641. 10.1038/s41592-022-01663-4.

40. Parikh, K., Antanaviciute, A., Fawkner-Corbett, D., Jagielowicz, M., Aulicino, A., Lagerholm, C., Davis, S., Kinchen, J., Chen, H.H., Alham, N.K., et al. (2019). Colonic epithelial cell diversity in health and inflammatory bowel disease. Nature 567, 49–55. 10.1038/s41586-019-0992-y.

41. Petukhov, V., Xu, R.J., Soldatov, R.A., Cadinu, P., Khodosevich, K., Moffitt, J.R., and Kharchenko, P.V. (2022). Cell segmentation in imaging-based spatial transcriptomics. Nat Biotechnol 40, 345–354. 10.1038/s41587-021-01044-w.

42. Pickert, G., Neufert, C., Leppkes, M., Zheng, Y., Wittkopf, N., Warntjen, M., Lehr, H.A., Hirth, S., Weigmann, B., Wirtz, S., et al. (2009). STAT3 links IL-22 signaling in intestinal epithelial cells to mucosal wound healing. J Exp Med 206, 1465–1472. 10.1084/jem.20082683.

43. Polanski, K., Young, M.D., Miao, Z., Meyer, K.B., Teichmann, S.A., and Park, J.E. (2020). BBKNN: fast batch alignment of single cell transcriptomes. Bioinformatics 36, 964–965. 10.1093/bioinformatics/btz625.

44. Ran, F.A., Hsu, P.D., Wright, J., Agarwala, V., Scott, D.A., and Zhang, F. (2013). Genome engineering using the CRISPR-Cas9 system. Nat Protoc 8, 2281–2308. 10.1038/nprot.2013.143.

45. Rao, D., Macias, E., Carbajal, S., Kiguchi, K., and DiGiovanni, J. (2015). Constitutive Stat3 activation alters behavior of hair follicle stem and progenitor cell populations. Mol Carcinog 54, 121–133. 10.1002/mc.22080.

46. Ritchie, M.E., Phipson, B., Wu, D., Hu, Y., Law, C.W., Shi, W., and Smyth, G.K. (2015). limma powers differential expression analyses for RNA-sequencing and microarray studies. Nucleic Acids Res 43, e47. 10.1093/nar/gkv007.

47. Schroeder, K.W., Tremaine, W.J., and Ilstrup, D.M. (1987). Coated oral 5-aminosalicylic acid therapy for mildly to moderately active ulcerative colitis. A randomized study. N Engl J Med 317, 1625–1629. 10.1056/NEJM198712243172603.

48. Schubert, M., Klinger, B., Klunemann, M., Sieber, A., Uhlitz, F., Sauer, S., Garnett, M.J., Bluthgen, N., and Saez-Rodriguez, J. (2018). Perturbation-response genes reveal signaling footprints in cancer gene expression. Nat Commun 9, 20. 10.1038/s41467-017-02391-6.

49. Shimokawa, M., Ohta, Y., Nishikori, S., Matano, M., Takano, A., Fujii, M., Date, S., Sugimoto, S., Kanai, T., and Sato, T. (2017). Visualization and targeting of LGR5(+) human colon cancer stem cells. Nature 545, 187–192. 10.1038/nature22081.

50. Smillie, C.S., Biton, M., Ordovas-Montanes, J., Sullivan, K.M., Burgin, G., Graham, D.B., Herbst, R.H., Rogel, N., Slyper, M., Waldman, J., et al. (2019). Intra– and Inter-cellular Rewiring of the Human Colon during Ulcerative Colitis. Cell 178, 714–730 e722. 10.1016/j.cell.2019.06.029.

51. Stoeckius, M., Zheng, S., Houck-Loomis, B., Hao, S., Yeung, B.Z., Mauck, W.M., 3rd, Smibert, P., and Satija, R. (2018). Cell Hashing with barcoded antibodies enables multiplexing and doublet detection for single cell genomics. Genome Biol 19, 224. 10.1186/s13059-018-1603-1.

52. Sugimoto, S., Ohta, Y., Fujii, M., Matano, M., Shimokawa, M., Nanki, K., Date, S., Nishikori, S., Nakazato, Y., Nakamura, T., et al. (2018). Reconstruction of the Human Colon Epithelium In Vivo. Cell Stem Cell 22, 171–176 e175. 10.1016/j.stem.2017.11.012.

53. Susaki, E.A., Tainaka, K., Perrin, D., Yukinaga, H., Kuno, A., and Ueda, H.R. (2015). Advanced CUBIC protocols for whole-brain and whole-body clearing and imaging. Nat Protoc 10, 1709–1727. 10.1038/nprot.2015.085.

54. Taniguchi, K., Wu, L.W., Grivennikov, S.I., de Jong, P.R., Lian, I., Yu, F.X., Wang, K., Ho, S.B., Boland, B.S., Chang, J.T., et al. (2015). A gp130-Src-YAP module links inflammation to epithelial regeneration. Nature 519, 57–62. 10.1038/nature14228.

55. Tetteh, P.W., Basak, O., Farin, H.F., Wiebrands, K., Kretzschmar, K., Begthel, H., van den Born, M., Korving, J., de Sauvage, F., van Es, J.H., et al. (2016). Replacement of Lost Lgr5-Positive Stem Cells through Plasticity of Their Enterocyte-Lineage Daughters. Cell Stem Cell 18, 203–213. 10.1016/j.stem.2016.01.001.

56. Tirosh, I., Izar, B., Prakadan, S.M., Wadsworth, M.H., 2nd, Treacy, D., Trombetta, J.J., Rotem, A., Rodman, C., Lian, C., Murphy, G., et al. (2016). Dissecting the multicellular ecosystem of metastatic melanoma by single-cell RNA-seq. Science 352, 189–196. 10.1126/science.aad0501.

57. Tomic, G., Morrissey, E., Kozar, S., Ben-Moshe, S., Hoyle, A., Azzarelli, R., Kemp, R., Chilamakuri, C.S.R., Itzkovitz, S., Philpott, A., and Winton, D.J. (2018). Phospho-regulation of ATOH1 Is Required for Plasticity of Secretory Progenitors and Tissue Regeneration. Cell Stem Cell 23, 436–443 e437. 10.1016/j.stem.2018.07.002.

58. van Es, J.H., Sato, T., van de Wetering, M., Lyubimova, A., Yee Nee, A.N., Gregorieff, A., Sasaki, N., Zeinstra, L., van den Born, M., Korving, J., et al. (2012). Dll1+ secretory progenitor cells revert to stem cells upon crypt damage. Nat Cell Biol 14, 1099–1104. 10.1038/ncb2581.

59. Wagner, D.E., and Klein, A.M. (2020). Lineage tracing meets single-cell omics: opportunities and challenges. Nat Rev Genet 21, 410–427. 10.1038/s41576-020-0223-2.

60. Wolf, F.A., Angerer, P., and Theis, F.J. (2018). SCANPY: large-scale single-cell gene expression data analysis. Genome Biol 19, 15. 10.1186/s13059-017-1382-0.

61. Wolf, F.A., Hamey, F.K., Plass, M., Solana, J., Dahlin, J.S., Gottgens, B., Rajewsky, N., Simon, L., and Theis, F.J. (2019). PAGA: graph abstraction reconciles clustering with trajectory inference through a topology preserving map of single cells. Genome Biol 20, 59. 10.1186/s13059-019-1663-x.

62. Wolock, S.L., Lopez, R., and Klein, A.M. (2019). Scrublet: Computational Identification of Cell Doublets in Single-Cell Transcriptomic Data. Cell Syst 8, 281–291 e289. 10.1016/j.cels.2018.11.005.

63. Ytterberg, S.R., Bhatt, D.L., Mikuls, T.R., Koch, G.G., Fleischmann, R., Rivas, J.L., Germino, R., Menon, S., Sun, Y., Wang, C., et al. (2022). Cardiovascular and Cancer Risk with Tofacitinib in Rheumatoid Arthritis. N Engl J Med 386, 316–326. 10.1056/NEJMoa2109927.

64. Yui, S., Azzolin, L., Maimets, M., Pedersen, M.T., Fordham, R.P., Hansen, S.L., Larsen, H.L., Guiu, J., Alves, M.R.P., Rundsten, C.F., et al. (2018). YAP/TAZ-Dependent Reprogramming of Colonic Epithelium Links ECM Remodeling to Tissue Regeneration. Cell Stem Cell 22, 35–49 e37. 10.1016/j.stem.2017.11.001.

65. Zheng, G.X., Terry, J.M., Belgrader, P., Ryvkin, P., Bent, Z.W., Wilson, R., Ziraldo, S.B., Wheeler, T.D., McDermott, G.P., Zhu, J., et al. (2017). Massively parallel digital transcriptional profiling of single cells. Nat Commun 8, 14049. 10.1038/ncomms14049.

